# Mesolimbic dopamine adapts the rate of learning from action

**DOI:** 10.1101/2021.05.31.446464

**Authors:** Luke T. Coddington, Sarah E. Lindo, Joshua T. Dudman

**Affiliations:** Howard Hughes Medical Institute, Janelia Research Campus, Ashburn, VA

## Abstract

Recent success in training artificial agents and robots derives from a combination of direct learning of behavioral policies and indirect learning via value functions. Policy learning and value learning employ distinct algorithms that optimize behavioral performance and reward prediction, respectively. In animals, behavioral learning and the role of mesolimbic dopamine signaling have been extensively evaluated with respect to reward prediction; however, to date there has been little consideration of how direct policy learning might inform our understanding. Here we used a comprehensive dataset of orofacial and body movements to understand how behavioral policies evolve as naive, head-restrained mice learned a trace conditioning paradigm. Individual differences in initial dopaminergic reward responses correlated with the emergence of learned behavioral policy, but not the emergence of putative value encoding for a predictive cue. Likewise, physiologically-calibrated manipulations of mesolimbic dopamine produced multiple effects inconsistent with value learning but predicted by a neural network-based model that used dopamine signals to set an adaptive rate, not an error signal, for behavioral policy learning. This work provides strong evidence that phasic dopamine activity can regulate direct learning of behavioral policies, expanding the explanatory power of reinforcement learning models for animal learning.

## Introduction

Biological and artificial agents learn how to optimize behavior through experience with an environment. Reinforcement learning (RL) theory describes the algorithms that allow an agent to iteratively improve its success through training^1^. Experience with the environment can be evaluated either by the success of an agent’s behavioral ‘policy’ that directly determines the actions performed (“policy learning”) or by an agent’s subjective expectations of reward that indirectly guide action (“value learning”). Over the last several decades much work has explored how mDA activity matches the predicted update signals (reward prediction errors, RPEs^2^) for value learning^3^. However, mDA activity also reflects a heterogeneous mix of signals and functions which may not be completely addressed by the predictions of value learning models^4–14^. Phasic mDA activity can be intertwined with the production and monitoring of action^6,10,11, 15–18^ and is determined at least in part by inputs from areas involved in determining behavioral policy^19,20^ . This calls for an exploration of whether and how broadening the scope of considered RL algorithms might inform our understanding of phasic mDA signals in biological agents.

Direct policy learning specifically offers untapped potential^21,22^ to provide “computational and mechanistic primitives”^23^ that explain dopamine functions, especially in the context of novel task acquisition by animals. First, direct policy learning methods have enjoyed substantial success in embodied learning problems in robotics that resemble problems faced by a behaving animal^24^. Second, under a wide set of conditions policy learning is the most parsimonious RL model that explains learned behavior^21^. Third, policy learning can be directly driven by behavioral performance error signals, in lieu of or in addition to, RPEs^25,26^, connecting them to diverse observations of learning in dopamine-recipient brain areas^10,27–35^. Finally, policy learning methods facilitate explicit modeling of meaningful variability^36,37^ in individual behavioral learning trajectories as a search through the space of policy parameterizations.

It can in fact be a criticism of policy learning that learning trajectories can be too variable; while conducive to modeling individual differences, this feature can produce suboptimal learning^38,39^. A powerful solution is to set an optimal update size for each trial according to some heuristic for how useful each trial could be for learning^40^. Doing so independently of the performance feedback that directs learning can enhance useful variability while suppressing noise^24,38,41^. Such “adaptive learning rates” have led to fundamental advances in machine learning^41,42^, and can also make models of animal learning more realistic^43,44^. Thus, insights from policy learning leads to an intriguing hypothesis for phasic mDA activity that has not, to date, been explored. Phasic mDA activity could be a useful adaptive learning rate signal, given its correlations to novel and salient stimuli, upcoming actions, and prediction errors, all of which are useful heuristics for identifying key learning moments during which learning rates should be elevated. Alternatively, mDA activity correlates with performance errors during avian song learning^28,45^, suggesting that in mammals it could also dictate error-based updates to behavioral policies - a role more analogous to conveying RPEs for value learning^45^. The establishment of policy learning models of canonical animal behavioral tasks is required to distinguish amongst these possibilities.

Here we develop a policy learning account of the acquisition of classical trace conditioning in which behavior is optimized to minimize the latency to collect reward once it is available, inspired by observations of this process in naive mice. A multidimensional dataset of behavioral changes during acquisition could be seen to drive improvements in reward collection performance, while a novel policy learning model quantitatively accounted for the diverse learned behavior of individual animals. mDA activity predicted by the component of this model which sets an adaptive learning rate closely matched fiber photometry recordings of mDA activity made continuously throughout learning. Individual differences in initial phasic mDA responses predicted learning outcome hundreds of trials later in a manner consistent with dopamine modulating learning rate. Optogenetic manipulation of ventral tegmental area (VTA) DA neurons was calibrated to physiological signals and triggered in closed-loop with behavior in order to provide a key test of the hypothesis that phasic mDA activity modulates learning rate as a distinct alternative to signaling signed errors. Together these results define a novel function for mesolimbic dopamine in adapting the learning rate of direct policy learning (see Logic Outline, Ext. Data Fig. 9).

## Results

### Individual differences in learning trajectories over acquisition of a cue-reward pairing

We tracked multiple features of behavioral responses to classical trace conditioning in thirsty mice that had been acclimated to head fixation but had received no other “shaping” or pre-training. Sweetened water reward was “cued” by a 0.5 s auditory cue (10 kHz tone) followed by a 1 s delay, except on a small number of randomly interleaved “uncued” probe trials (∼10% of total trials). While reward was delivered irrespective of behavior, mice still learned to optimize reward collection, as assayed by monotonic decreases in latency to collect reward across training (Fig. 1b-c). We measured multiple features of behavior to understand how idiosyncratic learning across individual mice subserved performance improvements: an accelerometer attached to the moveable basket under the mice summarized body movements^8^, while high resolution video was used to infer lick rate, whisking state, pupil diameter, and nose motion. We reasoned that reward collection performance could be improved along two dimensions: preparation for reward delivery and reaction to its sensory components (Fig. 1b-c). “Preparatory” behavior was assayed across lick, body, whisker, and pupil measurements as the total amount of activity during the delay period between cue and reward. “Reactive” behavior was assayed across nose, body, and whisker measurements as the latency to initiate following reward delivery.

**Figure 1.**
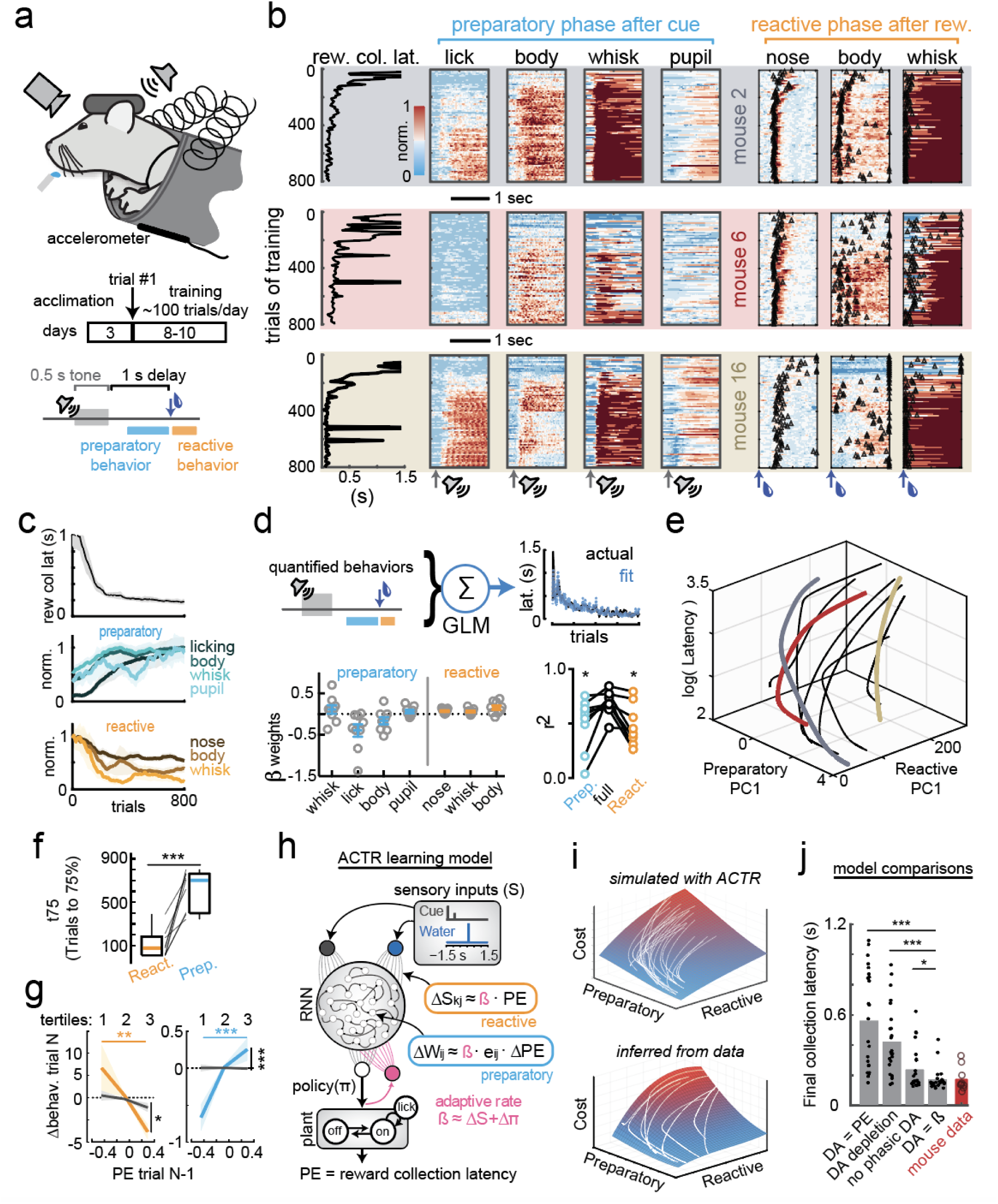
Changes to behavioral policy correlate with improved reward collection performance. **a)** Naive, thirsty, head-fixed mice underwent classical trace conditioning. **b)** Reward collection latency (leftmost column) compared to normalized heat maps of preparatory measures of licking, body movement, whisking probability and pupil diameter (middle 4 columns), and reactive measures of nose motion, body movements, and whisking probability (right 3 columns, with mean first response following reward delivery indicated by black triangles) for standard trials in which a 0.5 s auditory cue (grey arrows at cue start) predicted 3 μL sweetened water reward (blue arrows), averaged in 10-trial bins across training. Each color-coded row summarizes an example mouse’s learning trajectory, with the background color identifying the same mice in Fig 1e. **c)** 100 trial moving means of reward collection latency (top) across all 9 mice, compared to normalized preparatory (middle) and reactive (bottom) measures across 800 trials of training. **d)** (top)Preparatory and reactive measures as in panel (c) used as predictors of reward collection latency in a generalized linear model for each mouse. (bottom, left) Weights of each predictor in the GLM for each of 9 individual mice. (bottom, right) Preparatory predictors alone (blue) or reactive predictors alone (orange) provided worse predictions than the full model. **e)** Abstract learning trajectories were described as exponential fits to the first principal component of reactive and preparatory measurement variables, then plotted against the latency to collect reward for all mice (example mice from panel C: red, yellow, grey; all other mice: thin black), to visualize the relationship of inferred policy updates to reward collection performance. **f)** Number of trials to reach 75% of max learned performance for reactive (orange) and preparatory (blue) learning curves (thin gray lines: individual animals). p<0.001; ranksum test. **g)** Comparing the change in reactive (left, orange) and preparatory (right, blue) trajectory on each trial to the inferred performance error (PE) on the previous trial, overlaid with shuffled controls (black lines; compared to shuffle of trialwise PE for all other mice). Trials were binned into tertiles of PE across training. Significance testing with 2-way ANOVA; Tukey-Cramer post-hoc. *, p<0.05; **, p<0.01; ***, p<0.001. **h)** The ACTR model (see methods) learned a control policy (π) for the lick plant as the output from an RNN that received sensory inputs upon cue onset and offset (purple) and water reward delivery (red). Weights of connections between neurons were updated according to the ACTR learning rule summarized in the equation. S_kj_: weight of the connection between the k-th (tone or reward) sensory input and the output unit of RNN neuron. W_ij_: weight of the connection between the i-th neuron and the j-th neuron in RNN. *β*: adaptive learning rate set by a mDA-like signal from a feedback neuron (pink) summing the derivatives of sensory input (S) and policy output (π). E_ij_: eligibility trace for node perturbation at the synapse between the i-th neuron and the j-th neuron. PE: performance error derived from the latency to collect reward on the current trial. **i)** (top) Objective surface to visualize policy gradient calculated from ACTR model using arbitrary combinations of transient and sustained components (mean of 1000 simulations per point), overlaid with learning trajectories from individual initializations (white). (bottom) Objective surface fit (2^nd^ order polynomial surface) from observed mouse data, overlaid with observed learning trajectories (white, from panel (e)). **j)** Final collection latencies for different versions of ACTR model (grey bars, individuals as dots) compared to observed performance in mice (red bar, individuals as circles)

Preparatory and reactive components of learned behavior exhibited noisy, but monotonic trajectories on average (Fig 1c), with variable magnitudes and time courses across individuals (Fig. 1b). To assess whether changes in both preparatory and reactive components were correlated with improvements in reward collection performance, we built generalized linear models (GLM) to predict reward collection latency across training in each mouse (Fig. 1d). GLMs using preparatory and reactive behavioral measures as predictors captured much of the variance in reward collection efficiency over training (r^2^ = 0.69 ± 0.11; r^2^ with shuffled responses = 0.01 ± 3e^-4^). Each predictor’s weighting could vary widely from mouse to mouse, with preparatory licking having the most consistent relation to reward collection latency (Fig. 1d). However, both preparatory and reactive variables were necessary to most accurately predict reward collection latency (Fig. 1d, Friedman’s: P = 0.0003; preparatory only r^2^ = 0.51 ± 0.24, vs full model P = 0.004; reactive only r^2^ = 0.46 ± 0.20, vs full model P = 0.002).

The additive contribution of preparatory and reactive components to explained variance in performance and their significantly different time courses (Fig. 1f) suggest these two learning components are dissociable processes. Consistent with direct policy updates by a performance error related to reward collection latency, we found that updates to both the reactive and preparatory behavior on each current trial were significantly related to reward collection latency on the previous trial (Fig. 1g). Thus, we describe updates to the behavioral policy for each mouse as a trajectory through an abstract ‘learning space’ spanned by two components (preparation and reaction) that together explain improvements in reward collection performance (Fig. 1e).

### ACTR: a direct policy learning model of classical conditioning

The above data explain novel acquisition of novel trace conditioning as a problem of iteratively optimizing an effective control policy for reward collection. To formalize this concept, we first began by specifying a behavioral ‘plant’ (Ext. Data Fig. 1d) that captured the statistics of rodent licking behavior as a state model that transitions between quiescence and a licking state which emits a physiological lick frequency^46^. A continuous control policy (π(t)) determines the forward transition rate to active licking. The reverse transition rate reflects a bias towards quiescence that decreases in the presence of water such that licking is sustained until collection is complete. The control policy was learned as the additive combination of output from a recurrent neural network (RNN) modeling preparatory learning and a feedforward sensorimotor pathway modeling reactive learning (Fig. 1h; see Methods for model details and code).

Importantly, the optimal policy (identified by a search through the space of potential RNNs (Ext. Data Fig. 1a-b)) minimized performance cost by producing preparatory cued licking that depends upon sustained dynamics in the RNN output.

Performance errors used to train the model were proportional to the difference between the policy output at the time of reward delivery and the latency to collect water reward (see Methods). Behavioral learning data demonstrated that all mice exhibited relatively rapid and stable reactive learning present in both cued and uncued trials. To replicate these properties in the model, reactive learning was implemented as a change in the feedforward weights from sensory inputs to behavioral policy output updated in proportion to the performance error (Fig. 1h). Preparatory learning in the RNN was similarly proportional to performance errors, but used the change in performance error as is customary in policy learning algorithms^25^. The combination of reactive and preparatory learning robustly converged on stable, near optimal policies that led to dramatic reductions in the latency to collect reward (Fig. 1h-j). To stabilize policy across a range of model initializations, an adaptive learning rate for each trial was controlled by a feedback unit (pink output unit in Fig. 1h); activity of the feedback unit was the sum of the state change in the behavioral plant (akin to an efference copy of reward-related action initiation commands^47^) and the change in behavioral policy at the time of reward delivery (akin to a reward-predictive sensory evidence^8^). This feedback scheme has a direct and intentional parallel to the phasic activity of midbrain DA neurons in this task, which is well described as the sum of action- and sensory-related components of reward prediction^8,15^. Overall, our approach adds an adaptive rate component inspired by supervised learning optimization methods^38,41,48^ to a biologically plausible rule for training RNNs^49^ that itself drew inspiration from node perturbation methods^50^ and classic direct policy optimization methods known as ‘REINFORCE’^1,25^. Hence, we refer to the complete model as ACTR, for <u>A</u>daptive rate, <u>C</u>ost of performance to <u>R</u>EINFORCE.

ACTR successfully reproduced many significant aspects of mouse behavioral learning data. For repeated ACTR simulations (n=24) with a range of initializations, latency to collect rewards declined comparably to observed mouse behavior over training (Fig. 1j, including an equivalent cued performance gain (Ext. Data Fig. 1e)). Learning trajectories (Ext. Data Fig. 1c) and cost surfaces (Fig. 1i) calculated from a range of model initializations compared well qualitatively and quantitatively to those inferred from mouse data (Fig. 1i; see Methods). Finally, modeling the adaptive rate term (β) after phasic midbrain DA activity (see Fig 3f for comparison of modeled to actual) was well supported by comparing end performance of the full model to modified versions (Fig. 1j) in which: (1) the DA-like feedback unit signaled performance error instead of rate (Sup Fig 1f; significantly worse performance, ranksum p<2x10^-7^); (2) learning rate was globally reduced (akin to DA depletion^51^; significantly worse performance, ranksum p<2x10^-6^)); (3) a basal learning rate was intact but there was no adaptive component (akin to disruption of phasic mDA activity^52^; significantly worse performance, ranksum p=0.02)). Thus initial trace conditioning is well described as the optimization of reward collection behavior, and best approximated when DA-like signals act not as signed errors directing changes to the policy, but instead adapting the size of the learned update on each trial.

### Mesolimbic dopamine signals correlate with behavioral learning trajectory

We measured mDA activity in the above mice, which were DAT-Cre::ai32 mice that transgenically expressed Chr2 under control of the dopamine transporter promoter, by injecting a Cre-dependent jRCaMP1b virus across the midbrain (Fig. 2a, b) ^8,53^. This combined optogenetic/fiber photometry strategy also allowed for calibrated DA manipulations in later experiments. Optical fibers were implanted bilaterally over the VTA, and unilaterally in the nucleus accumbens core (NAc), and in the dorsal medial striatum (DS) (Fig. 2a). We recorded signals from the NAc in all mice, with some additional simultaneous recordings from ipsilateral VTA (n=3) or contralateral DS (n = 6). NAc-DA reward responses became better aligned to reward delivery across training but did not decrease significantly (trials 1-100: 0.82 ± 0.21 z, trials 700-800: 1.16 ± 0.23 z, signed-rank p = 0.13), even as cue responses steadily increased (Ext. Data Fig. 2a). These dynamics recapitulated our previous observations from somatic activity in the VTA and SNc, and indeed closely resembled simultaneously recorded VTA-DA signals Ext. Data Fig. 2a-e). In contrast, DS-DA developed cue and reward responses only upon further training, matching previous reports^11,54,55^ (Ext. Data Fig. 2a-e), and indicating that mesolimbic (e.g. VTA→NAc) reward signals are of specific interest during the initial learning period studied here.

**Figure 2.**
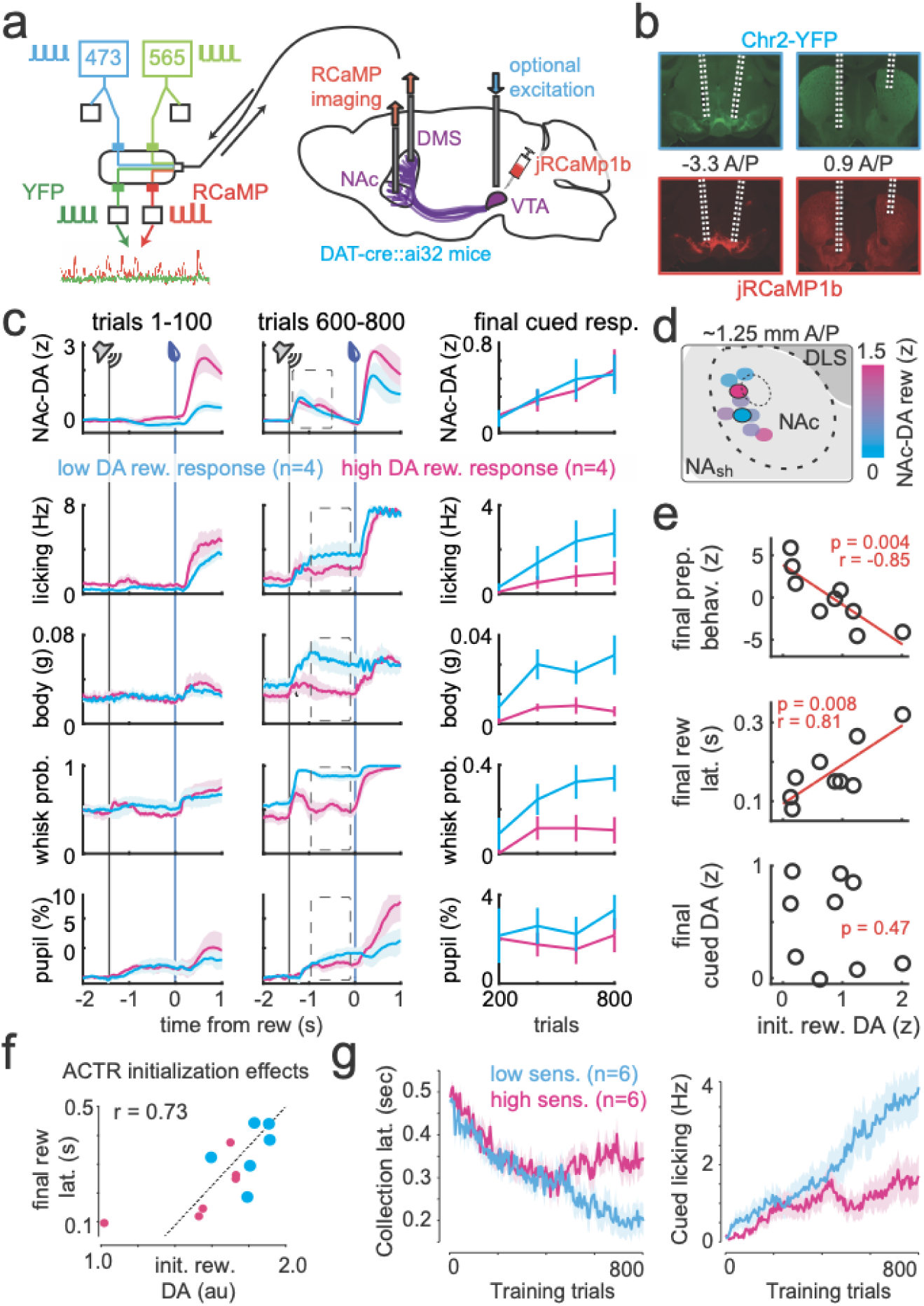
Individual differences in mesolimbic DA signals correlate with learned behavioral policy. **a)** (left) Fiber photometry hardware schematic. 10% duty cycle 473 and 656 nm excitations were offset from eachother and split between the main filter cube and photodectors (white squares) that measure output power. Excitation and emission of eYFP and jRCaMP1b fluorescence were conveyed by one cable between the filter cube and the brains of head-fixed animals. YFP and RCaMP emissions were measured at separate filter cube outputs. (right) jRCaMP1b was virally expressed bilaterally in the VTA and SNc of DAT-cre::ai32 mice, allowing measurement and the option for simultaneous manipulation of mesostriatal DA circuits. **b)** Histology showing example fiber paths and virus expression. **c)** (left) NAc-DA, licking, body movements, whisking probability, and pupil diameter measurements for the mean of the 4 animals with lowest (blue) and highest (pink) NAc-DA reward signals over the initial 100 trials, shown for trials 1-100 compared to trials 600-800. (right) Means of responses in the analysis windows indicated at left (dashed gray boxes) across training. **d)** Visualization of fiber locations for each mouse (n=9), color-coded according to the size of their initial NAc-DA reward signals. **e)** Correlations of initial NAc-DA reward responses (trials 1-100) with final combined preparatory behavior magnitude (see Methods)(top), final latency to collect reward (middle), and lack of correlation to final cued NAc-DA response (bottom) (n=9 mice). **f)** Simulations with low (small dots, n=6) or high (large dots, n=6) initial reward-related sensory input exhibited a significant correlation between initial (trials 1-100) predicted mDA reward response and final reward collection latency. **g)** Reward collection latency (left) and preparatory cued licking (right) for simulations with low (cyan) and high (magenta) initial reward-related sensory input.

We thus proceeded to examine how individual differences in mesolimbic reward signals were related to the individual differences in behavioral learning detailed in Figure 1. We found substantial inter-animal variance in initial NAc-DA responses in the first 100 trials that was not related to anatomical location of fibers (Fig. 2d; initial NAc-DA reward: A/P: p=0.5, M/L: p=0.4, D/V: p=0.5; multiple linear regression all axes, p = 0.7). Surprisingly, initial NAc-DA reward signals were negatively correlated with the amount of preparatory behavior at the end of training (Fig. 2c, e; NAc-DA reward_trials 1-100_ vs preparatory index_trials 700-800_, r = -0.85, p = 0.004), as well as the speed of reward collection (Fig. 2e; NAc-DA reward_trials 1-100_ vs reward collection latency_trials700-800_, r = 0.81, p = 0.008). This relationship was specific for preparatory behaviors (Ext. Data Fig. 2g; NAc-DA reward_trials 1-100_ vs reactive index_trials 700-800_, p = 0.24), and was robust as each mouse’s initial dopamine reward signals could be accurately predicted from quantifications of final preparatory behavior (Ext. Data Fig. 2f; actual vs predicted r = 0.99, p < 0.0001).

The negative correlation between DA reward signal and behavioral learning is not consistent with the magnitude of phasic mDA activity determining or correlating with the error used for a learning update. However, it is potentially consistent with phasic mDA activity reflecting action-related and sensory-related components of the control policy. At initialization of the ACTR model, no preparatory actions have been learned so the DA signal is dominated by the initial reactive response to sensory input at reward delivery. Adjusting the strength of this sensory input at model initialization scales the initial DA reward response magnitudes similarly to the range observed in mice (Fig. 2f). Intriguingly, these initialization differences in ACTR simulations predicted a delayed collection latency at the end of training (Fig. 2g; r=0.73, p=0.007) due to a reduced development of a preparatory licking policy (Fig. 2g), mirroring our results in mice and demonstrating that stronger reactive responses to sensory information can impair the learned development of preparatory responses (and thus ultimately impair performance). Thus the ACTR model provides the insight that the initial strength of the feedforward sensorimotor component that is reported by mDA signals can reflect the state of the network at task initiation, resulting in significant difference in the course of learning.

### Only large dopamine manipulations drive value-like learning

Work exploring direct roles of DA in movement^11,56^ or motivation^57,58^ suggests that phasic cue responses provoke or invigorate preparatory behavior. Indeed, learned NAc-DA cue responses were correlated with cued licking across mice (Ext. Data Fig. 3c). However, at the end of regular training some mice experienced an extra session in which VTA-DA stimulation was triggered on cue presentation on a random subset of trials (Ext. Data Fig. 3). Increasing mesolimbic cue responses in this way had no effect on cued licking in the concurrent trial (control 2.3 ± 1.1 Hz, stim. 2.3 ± 1.0 Hz, p > 0.99). Thus within this context (though not necessarily others^59^), the magnitude of NAc-DA cue signals only correlates with learned changes in behavioral policy but does not appear to directly regulate preparatory behavior in anticipation of reward delivery^8,60^.

Surprisingly, individual differences in initial NAc-DA reward signals were not correlated with the learning of NAc-DA cue signals (reward_trials 1-100_ vs. cue_trials 700-800_, p = 0.5, Fig 2e). This could argue that DA reward signals are not a driving force in the learning of cue signals. This is surprising given that results in rodents^60,61^ and monkeys^62^ provide specific evidence for value learning effects following exogenous VTA-DA stimulation. However, reward-related DA neuron bursting is quite brief (<=0.2 s) in our task^8^ as well as in canonical results across species^3^, raising the question of whether high power, longer duration stimulation recruits the same learning mechanisms as briefer, smaller reward-sized responses. We next used our ability to simultaneously manipulate and measure mesolimbic DA^53^ to examine the function of brief DA transients calibrated to match reward responses from our task.

Following initial training on the trace conditioning paradigm, we introduced mice to a novel predictive cue - a 500-ms flash of light directed at the chamber wall in front of the mouse. After 10 introductory trials, this visible cue stimulus was paired with exogenous VTA-DA stimulation after 1 s delay for 5 daily sessions (∼150 trials per session). One group of randomly selected mice received VTA-DA stimulation calibrated to uncued reward responses (150 ms at 30 Hz and 1-3 mW steady-state power, stim response = 1.4 ± 0.3 uncued reward response, n=10), while the complement received larger, uncalibrated stimulations (500 ms at 30 Hz and 10 mW steady-state power, stim response = 5.5 ± 0.8 uncued reward response) (Fig. 3a). After 5 sessions, we found that the group receiving calibrated, reward-sized stimulation did not exhibit NAc-DA cue responses above baseline (0.0 ± 0.2 z, p = 0.8), whereas the large, uncalibrated stimulation group exhibited substantial NAc-DA cue responses (0.5 ± 0.2 z, p = 0.02) (Fig 3b).

**Figure 3.**
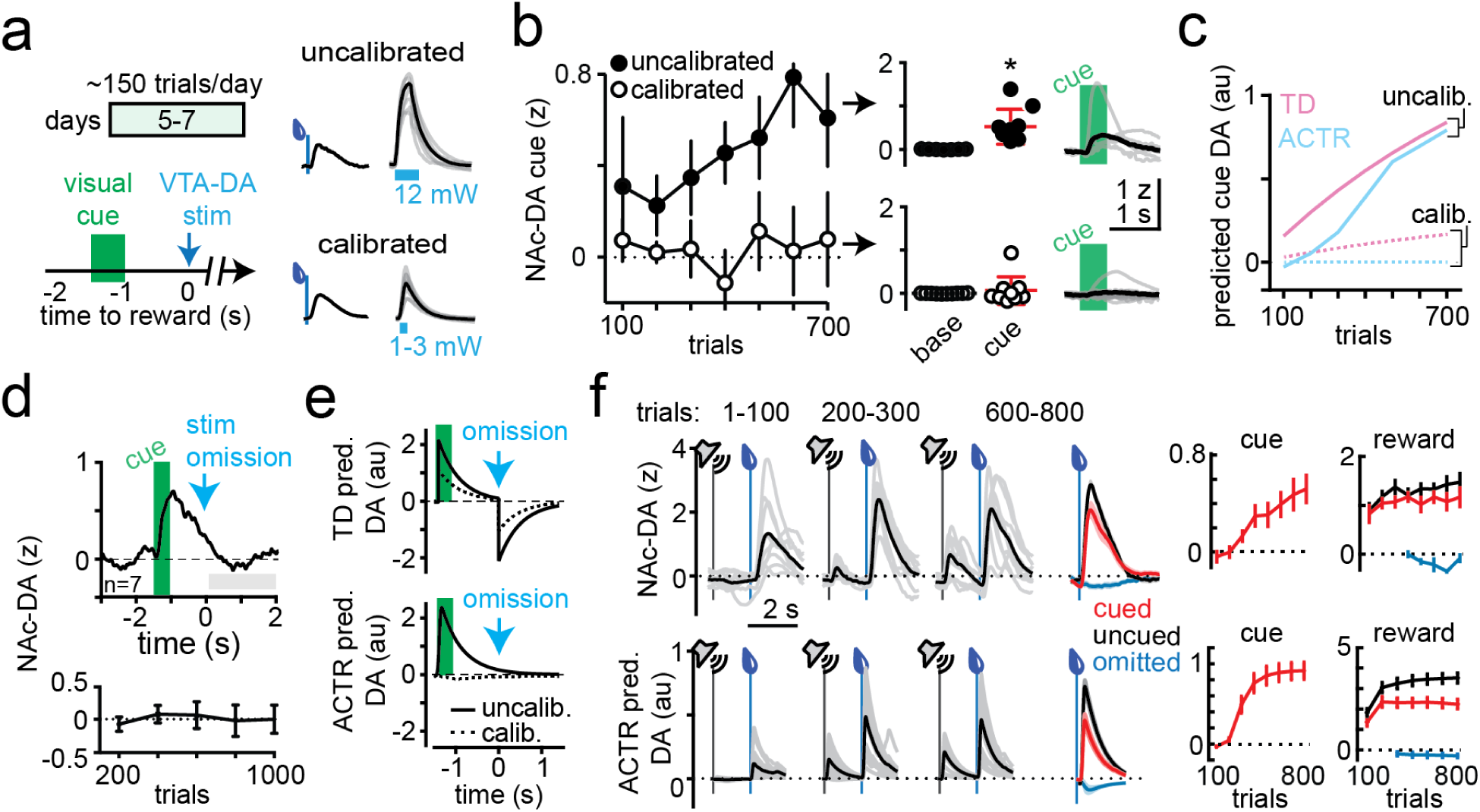
Large mesolimbic DA manipulations drive value-like learning. **a)** (left) Experimental design for VTA-DA stimulation predicted by a 0.5 s light flash at the front of the behavioral chamber. (right) Mean uncued NAc-DA reward responses vs VTA-DA stimulation responses (right, individuals in gray, mean in black) for mice that either received large, uncalibrated stimulation (top; 30 Hz, 12 mW for 500 ms) or stimulation calibrated to reward responses (bottom; 30 Hz, 1-3 mW for 150 ms). **b)** (left) jRCaMP1b NAc-DA cue responses across training for mice that received large stimulations (5x the size of reward responses; filled circles) or calibrated stimulations (1x the size of reward responses; open circles). (right) Quantified and raw data for mean NAc-DA traces after 750 training trials (5 sessions) with uncalibrated (top) and calibrated (bottom) stimulation. **c)** Predicted DA cue responses for the experiment in (a) simulated with a Temporal Discounting (TD) value learning model (light pink) vs ACTR (light blue) **d)** (top) Mean NAc-DA signal after 7 sessions of training with uncalibrated stimulation, on probe trials with omitted stimulation that were delivered 10% of the time, with quantification over training (bottom) **e)** Predicted DA responses by TD (top) and ACTR (bottom) models, for uncalibrated (bold line) and calibrated (dotted line) DA stimulation **f)** (top) Measured NAc-DA responses across training, including mean responses to reward on cued (red), uncued (black), and omission (blue, aligned to point at which reward should have been delivered) trials, with quantification over training (right) for cue and reward responses. (bottom) Same as top except for predicted DA responses in the ACTR model.

The emergence of cue signals following uncalibrated dopamine stimulation was captured in ACTR by introducing a nonlinearity in which larger, more sustained DA activation was modeled as a large modulation of learning rate coupled with a change in performance error encoding (Fig 3c-e; see Methods). This coupled effect in the model enhanced cue encoding in a manner similar to the predictions of value learning; however, it was also distinct in that such change in the encoding of cues was not accompanied by the suppression of DA activity on omission of the laser stimulus expected from value-learning models (Fig. 3d, e). In contrast to prior observations of inhibition following omission of expected stimulation in the context of consistent, overt behavioral responses to cues^18,63^, only a brief bout of body movement accompanied cue learning in the current paradigm (Ext. Data Fig. 4). This further suggests that ongoing action control may be fundamental to the observation of omission-related inhibition^8,15,18^.

In separate experiments, calibrated and uncalibrated VTA-DA stimulations had similar input-output properties across the medial prefrontal cortex and the dorsal-to-ventral axis of the striatum, suggesting that the recruitment of value-like cue learning by uncalibrated stimulation was related to the magnitude or duration of the uncalibrated signal rather than an increased spatial spread (Ext. Data Fig. 5). Together these data provide further evidence that the magnitude of phasic mDA reward responses measured in our task are not sufficient to drive value-like learning of predictive cue responses, even while larger stimulation flooding the same downstream regions with higher DA concentrations are sufficient to teach phasic responses to a cue that predicts DA stimulation.

### DA activity correlates with RPE but functions as an adaptive learning rate

We next elaborate on the role of DA reward signals in performance-driven direct policy learning. NAc-DA signals predicted by ACTR can be visualized by convolving its DA-like adaptive learning rate signal with a temporal kernel matched to the kinetics of jRCaMP1b^64^.

This predicted phasic DA photometry signal corresponded closely to experimentally measured NAc- and VTA-DA activity across training (Fig. 3f, Ext. Data Fig. 2a). Notably, modeling DA signals as the sum of action and sensory components in a control policy reproduces canonical RPE correlates, despite the lack of an explicit RPE computation (Fig. 3f, Ext. Data Fig. 6). It also predicts that DA reward signals should reflect the evolution of reward collection policy across learning. While animals’ policies are not directly observable, the presence or absence of preparatory licking on a given trial of behavior are a noisy correlate of differences in underlying behavioral policy. Indeed, in both ACTR and mouse data, differential reward responses on trials with (‘lick+’) or without (lick-) preparatory licking emerged in correlation with the extent of learning (Ext. Data Fig. 7).

While the differential DA dynamics on lick- vs lick+ trials may be expected from other models, a potential distinguishing characteristic is the way in which these differential signals are predicted to affect learning. ACTR (or in general any policy learning model^25^) evaluates whether specific changes in parameter values lead to relatively better or worse performance on a given trial. The mDA-like signal (*β*) adaptively determines learning rate, *i.e. how fast* to change. The actual *sign* of the change (towards or away from the new policy parameterization) is determined independently of mDA by the performance error (PE). For example, on trials in which collection latencies get longer relative to recent performance, the sign of the PE directs away from the current policy parameterization. Enhancing the phasic mDA-like signal at reward (increasing the learning rate) on such a trial would then bias *away* from that policy in the future (resulting in “less” of the associated behavior).

This exact relationship is present on average, though somewhat counterintuitively, for lick- and lick+ trials as training progresses. The stochasticity of the licking plant ensures that some trials with a good policy (lick+ trials with preparatory licking) can result in worse collection latencies relative to recent trials, *i.e.* negative performance errors. This could occur for instance on a lick+ trials in which preparatory licking suboptimally terminates prior to reward delivery. Maximizing learning rate on lick+ trials with negative performance error should bias away from a policy of preparatory licking, while maximizing learning rate on good lick+ trials should only bias towards preparatory licking by a much smaller amount due to vanishing errors as learning progresses. Enhancing learning rates on lick+ trials would thus, in aggregate, bias *away* from those types of trials occurring later in learning. The converse is true for lick- trials, where amplification of negative performance errors should bias towards preparatory licking (away from policies with no preparatory behavior).

These paradoxical effects of enhanced rates impairing learning can be demonstrated in ACTR simulations, as trial-type dependent enhancement of learning rate indeed produced opposite signed effects on preparatory licking behavior (Fig. 4g). Thus the larger NAc-DA reward signals observed for lick- trials in mice (Ext. Data Fig. 7c, d) is not only predicted by our direct policy learning model but may contribute to improved performance by increasing preparatory licking. Importantly, this dissociation in predicted effects of DA stimulation dependent upon trial type is unique to modulation of learning rates. When DA reward signals are modeled as signed errors - PEs in ACTR or analogously RPEs in a value-learning model - stimulation is predicted to enhance cue encoding and preparatory licking independent of trial-type (Fig. 4g).

**Figure 4.**
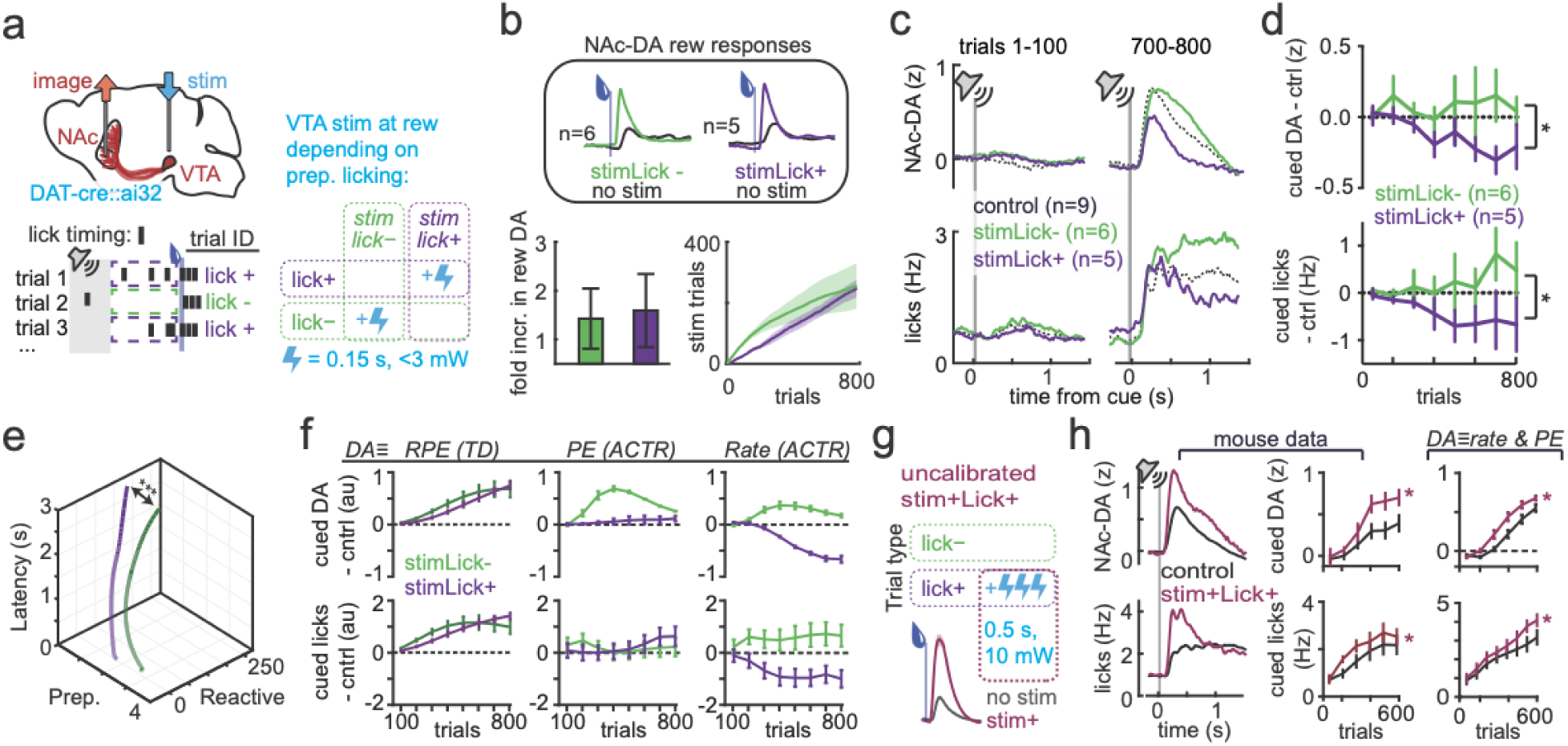
Role of mesolimbic DA in learning is consistent with an adaptive policy learning rate. **a)** (top) Simultaneous measurement and manipulation of mesolimbic DA signals. (bottom) Closed-loop experiment design, where different groups of mice received VTA stim (0.15 s, 30 Hz) concurrent with reward delivery on either lick- trials (“stim lick-”) or lick+ trials (“stime lick+). **b)** (top) Mean NAc-DA reward responses across training with (colored traces) and without (black traces) exogenous stimulation, for stimLick- (green)and stimLick+ (purple) mice. (bottom) Fold increase in NAc-DA reward signals on stimulated trials and cumulative sum of stimulated trials across training for stimLick- (green) and stimLick+ (purple). **c)** NAc-DA (top) and licking (bottom) during early (trials 1-100, left) and late (trials 700-800, right) training for control (black), stimLick- (green) and stimLick+ (purple) animals. **d)** NAc-DA cue responses (top) and cued licking (bottom) for StimLick- (green) and stimLick+ (purple) mice across training, displayed as the difference from control mice. **e)** 3d learning trajectories as in Fig. 1e, comparing preparatory and reactive components of behavior to the latency to collect reward across training. **f)** Predicted DA cue responses (top) and cued licking (bottom) compared for 3 possible functions of phasic DA signals: acting as the RPE signal in TD value learning model (left), biasing the performance error term (PE) in the ACTR policy learning model (middle), or setting the adaptive rate term (β) in ACTR (right). **g)** (top) Experimental design for new group of mice that experienced a “stim+Lick+” contingency: they received large, uncalibrated VTA-DA stimulation on lick+ trials. (bottom) NAc-DA reward responses on stimulated (magenta) and unstimulated (gray) trials **h)** NAc-DA cue responses (top) and cued licking (bottom) for control (black) and stim+Lick+ (light purple) mice.

We thus performed the same experiment in mice, selectively increasing DA reward signals through optogenetic stimulation in the VTA contingent upon preparatory cued behavior. Separate groups of animals experienced each of the following stimulation contingencies: “stimLick+” animals received VTA-DA stimulation at the moment of reward delivery on trials in which we detected licking in the 750 ms preceding reward delivery, while “stimLick−” animals received the same stimulation on trials in which no licking was detected during the delay interval (Fig. 4a). Crucially, stimulation was brief (150 ms) and calibrated to endogenous fiber photometry signals in each mouse to approximately double the endogenous NAc-DA reward response (Fig. 4b, see Methods,^8,53^). In order to account for the large discrepancy in stimulated trials that would arise between the two stimulation groups due to eventual predominance of lick+ trials, stimLick+ animals were limited to having a max of 50% of total trials stimulated in a given session. This resulted in a comparable number of stimulated trials between the two groups by the end of the training period (Fig. 4b).

Calibrated enhancement of reward-related activity in VTA→NAc-DA projections in this way had opposite effects on emerging delay-period behavior across the two stimLick contingencies (Fig. 4b-e). As in the ACTR model, behavior was biased in opposite directions for each contingency, with stimLick+ animals exhibiting lower and stimLick− animals exhibiting higher preparatory licking ((trials 600-800, stimLick+ 1.0 ± 0.7, stimLick- 0.6 ± 0.1, ANOVA F _1,72_ = 10.5, p = 0.002)). Furthermore, NAc-DA cue signals were biased in matching directions, with the stimLick− group also exhibiting higher NAc-DA cue responses vs stimLick+ (trials 600-800, stimLick+ 0.3 ± 0.1 z, stimLick- 2.6 ± 0.7 z, ANOVA F _1,72_ = 10.1, p = 0.002). Baseline licking examined just before trials began across training showed no correlation with the extent of learning (p=0.9) or initial NAc-DA magnitude (p=0.8), confirming that preparatory licking learning was indeed largely driven by the predictive cue (Ext. Data. Fig. 8).

The qualitative differences in effects of calibrated and uncalibrated stimulations (Fig. 3) suggests that uncalibrated stimulation could paradoxically reverse the effect on suppression of cued licking seen in the stimLick+ condition above. To test this possibility we repeated the stimLick+ experiment with a new set of mice, but this time augmented rewards on lick+ trials with large, uncalibrated VTA-DA stimulation (500 ms, at 30 Hz and ∼10 mW power)(Fig. 4h-i). Indeed, with this new larger exogenous stimulation, the stimLick+ contingency now resulted in *increased* NAc-DA cue responses within 600 trials (2-way ANOVA, stim group F_1,66_, p = 0.001) as well as *increased* cued licking (2-way ANOVA, stim group F_1,60_, p =0.01), reversing the sign of the effects of calibrated stimLick+ stimulation (Fig. 4d). These effects were well predicted by a modified version of the ACTR model in which large DA stimulations biased towards positive PE in addition to modulating learning rate (Fig. 4i, right), exactly as in the previous experiment in which large uncalibrated stimulation caused the emergence of NAc-DA responses to a predictive cue (Fig. 3a-c).

## Discussion

Here we used the core computational elements of machine learning–cost functions, circuit architecture, and learning rules^40,65^–to describe a novel function for midbrain dopamine in regulating a canonical animal learning paradigm, classical trace conditioning (see Logic Outline, Ext. Data Fig. 9). The discovery that the phasic activity of mDA neurons in several species correlated with a key quantity (RPE) in value learning algorithms has been a dramatic and important advance suggesting that the brain may implement analogous processes^2,3,66^. At the same time, reinforcement learning constitutes a large family of algorithms that include learning about not only expected values of environmental states, but also directly learning parameterized policies for behavioral control. Here we develop a novel and biologically plausible network-based formulation of policy learning that is consistent with many aspects of individual behavioral trajectories, but also closely matches observed mDA neuron activity during naive learning. Reinforcement learning in other situations in which anticipatory, approach, or operant behavior can be shown to minimize performance errors may also be explained as a descent along a policy gradient driven by performance evaluation.

Independent of the specific reinforcement learning algorithm, our analyses and experiments importantly discriminate between two potential functions of DA: a signed error term that governs the direction of learned changes and an unsigned rate term that governs how much of the error is captured by each update. The realization that DA activity is consistent with a role in modulating learning rate as opposed to signaling an error predicted that stimulation of mDA neurons could slow behavioral learning in some contingencies. Such a result would be paradoxical if DA activity functioned as an error. We note that stimulation of mDA neurons independent of behavioral performance, as routinely done, fails to distinguish these possibilities and thus a new experiment was required - calibrated manipulation of mDA activity in closed-loop with behavior (inferred policy states). We found a remarkable agreement between policy learning model-based predictions and experimental observations. Intriguingly, in separate experiments we discovered that uncalibrated mDA stimulation that was 3-5 times stronger than endogenous mDA activity (though with parameters similar to prior work, e.g.^61,63,67^) also can bias errors in addition to modulating learning rate. This suggests that dopamine-dependent attribution of motivational value to cues^61,63,68,69^ is at least partially dissociable from the regulation of policy learning rate within the same mesolimbic circuits. Such parallel functions could be complimentary, intriguingly mirroring the system of parallel policy- and value-learning networks implemented in AlphaGo^70^, one of the landmark achievements in modern artificial intelligence.

Value-like error signaling following higher power, longer simulations may depend on specific receptor recruitment within a circuit^71^ (as suggested by similar input/output relationships across tested regions (Sup Fig 5)), and/or differential recruitment of diverse^7,72,73^ dopaminergic circuits. A similar gain of function could occur when drugs of abuse enhance DA signaling, previously proposed to bias towards inappropriate positive learning errors^74^. Our data indicate that both significantly enhanced DA signaling and oversensitivity to sensory input can bias toward value-like learning that leads an animal to exhibit excessively strong reactive responses to cues at the expense of adaptive preparatory responses to predictive cues. This is perhaps akin to the excessive acquired responsiveness and/or innate sensitivity to drug-predictive cues thought to underlie the development of addiction^75^, and connects our results to previous observations of correlations between the magnitude of phasic DA signaling and individual differences in reward-related behaviors^69^. This suggests that policy learning, and specifically the reactive component in our ACTR model, may be a useful way to model the acquisition and expression of “incentive salience”^76^, although in our context phasic DA signals could only be shown to modulate learning, and not the expression of incentive salience on the current trial (Ext. Data Fig 3). Our results also underscore the importance of matching exogenous optogenetic manipulations to measured physiological signals, and support the idea that extended, high-magnitude mDA stimulation is an important model of addictive learning^77^ that can be dissociated from adaptive reinforcement learning processes.

There are many opportunities to extend the current model formulation to capture more biological reality and evaluate the biologically plausible, but currently incompletely tested, cellular and circuit mechanisms of posited ACTR learning rules. There is substantial prior evidence for the capacity of mDA activity to capture eligibility traces and modulate synaptic plasticity^78–80;^ however, our behavioral data and modeling call for further examination of multiple related learning rules governing reactive and preparatory learning. Given that adaptive control over the magnitude of learning rate can be a key determinant of machine learning performance in deep^40,41^ and recurrent neural networks^48^, studying how adaptive control of learning rates are implemented in animals, and especially across diverse tasks, may provide additional algorithmic insights to those developed here. Recent evidence also suggests that other neuromodulators in the brain may play distinct, putatively complementary roles in controlling the rate of learning^81^. Here we effectively identify a key heuristic apparent in phasic mDA activity that adapts learning rates to produce more stable and performant learning; however, we focused on a single behavioral learning paradigm and dopamine is known to be critical for a broad range of putative behavioral policies. Our work provides a perspective for future work to expand upon and identify other aspects of mDA activity that may be critical for the adaptive control of learning from action more broadly.

## Methods

### Animals

All procedures and animal handling were performed in strict accordance with protocols (11-39) that were approved by the Institutional Animal Care and Use Committee (IACUC) and consistent with the standards set forth by the Association for Assessment and Accreditation of Laboratory Animal Care (AALAC). For behavior and juxtacellular recordings we used 24 adult DAT-Cre::ai32 mice (3-9 months old) resulting from the cross of DAT^IRES*cre*^ (The Jackson Laboratory stock 006660) and Ai32 (The Jackson Laboratory stock 012569) lines of mice, such that a Chr2/EYFP fusion protein was expressed under control of the endogenous dopamine transporter Slc6a3 locus to specifically label dopaminergic neurons. Animals were housed on a 12-hour dark/light cycle (8am-8pm) and recording sessions were all done between 9am-3pm. Following at least 4 days recovery from headcap implantation surgery, animals’ water consumption was restricted to 1.2 mL per day for at least 3 days before training. Mice underwent daily health checks, and water restriction was eased if mice fell below 75% of their original body weight.

### Behavioral training

Mice were habituated to head fixation in a separate area from the recording rig in multiple sessions of increasing length over >= 3 days. During this time they received some manual water administration through a syringe. Mice were then habituated to head fixation while resting in a spring-suspended basket in the recording rig for at least two 30+ minute sessions before training commenced. No liquid rewards were administered during this recording rig acclimation, thus trial 1 in the data represents the first time naive mice received the liquid water reward in the training environment. The reward consisted of 3 μL of water sweetened with the non-caloric sweetener saccharin delivered through a lick port under control of a solenoid. A 0.5 s, 10 kHz tone preceded reward delivery by 1.5 s on “cued” trials, while 10% of randomly selected rewards were “uncued”. Matching our previous training schedule^8^, after three sessions, mice also experienced “omission” probe trials, in which the cue was delivered by not followed by reward, on 10% of randomly selected trials. Intertrial intervals were chosen from randomly permuted exponential distribution with a mean of ∼25 seconds. Ambient room noise was 50-55 dB, while an audible click of ∼53 dB attended solenoid opening upon water delivery and the predictive tone was ∼65 dB loud. Mice experienced 100 trials per session and one session per day for 8-10 days. In previous pilot experiments, it was observed that at similar intertrial intervals, behavioral responses to cues and rewards began to decrease in some mice at 150-200 trials. Thus the 100 trial/session limit was chosen to ensure homogeneity in motivated engagement across the dataset.

Some animals received optogenetic stimulation of VTA-DA neurons concurrent with reward delivery, contingent on their behavior during the delay period (see technical details below). Following trace conditioning with or without exogenous DA stimulation, 5 mice experienced an extra session during which VTA-DA neurons were optogenetically stimulated concurrently with cue presentation (Sup. Fig. 3). Mice were then randomly assigned to groups for a new experiment in which a light cue predicted VTA-DA stimulation with no concurrent liquid water reward (5-7 days, 150-200 trials per day). The light cue consisted of a 500 ms flash of a blue LED directed at the wall in front of head fixation. Intertrial intervals were chosen from randomly permuted exponential distributions with a mean of ∼13 seconds. Supplementary Table 1 lists the experimental groups each mouse was assigned to in the order in which experiments were experienced.

**Supplementary Table 1.**
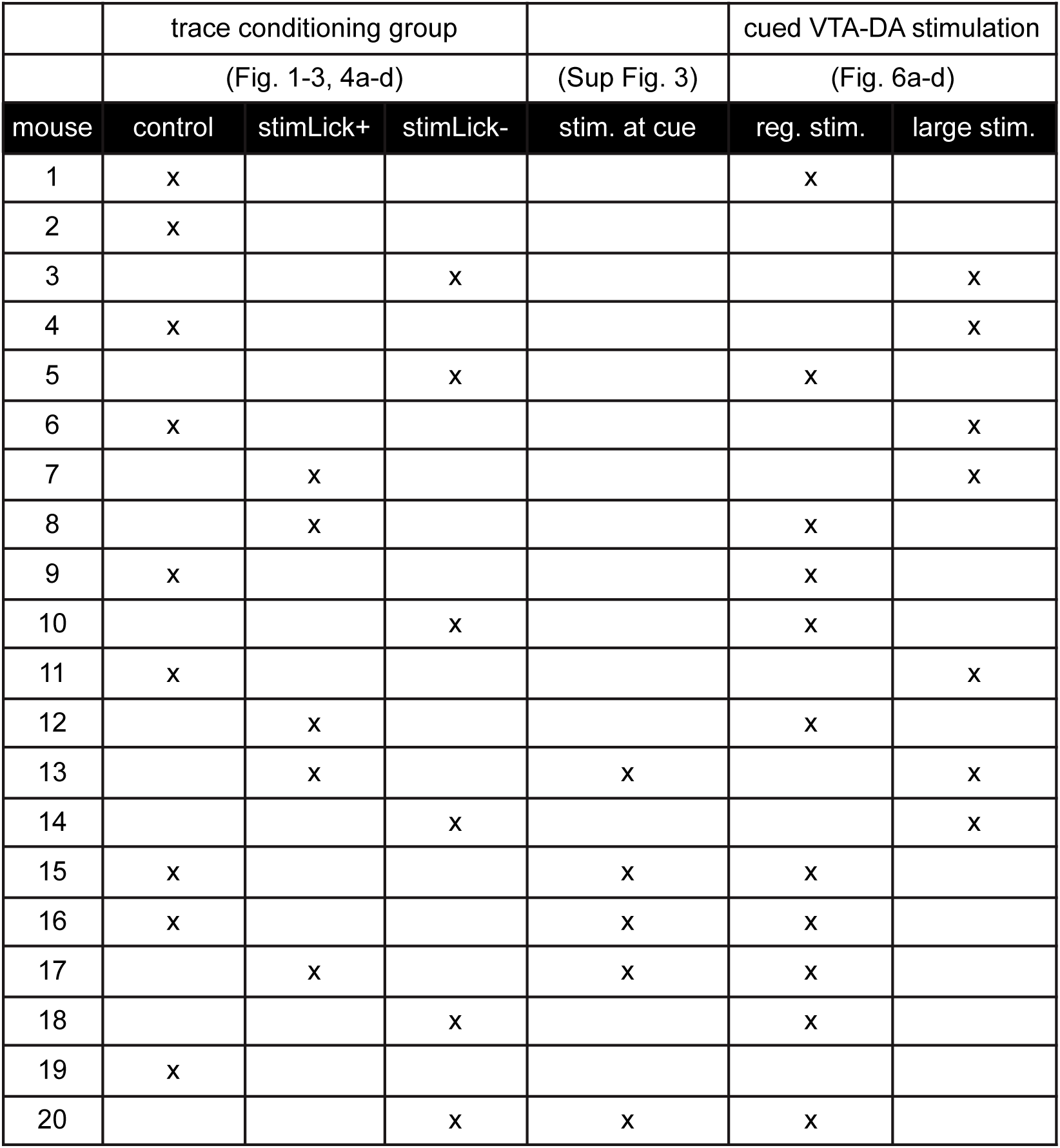
Successive experimental group membership for each mouse. Mice #21-24 are not included above as they were only used for a single experiment, the stim+Lick+ experiments in Fig. 4g-h

### Video and behavioral measurement

Face video was captured at 100 Hz continuously across each session with a single camera (Flea 3, FLIR) positioned level with the point of head fixation, at a ∼30° angle from horizontal. Dim visible light was maintained in the rig so that pupils were not overly dilated, while an infrared LED (model#) trained at the face provided illumination for video capture. Video was post-processed with custom matlab code (available at: www.github.com/).

Briefly, for each session, a rectangular region of interest (ROI) for each measurement was defined from the mean of 500 randomly drawn frames. Pupil diameter was estimated as the mean of the major and minor axis of the object detected with the MATLAB ‘regionprops’ function, following noise removal by thresholding the image to separate light and dark pixels, then applying a circular averaging filter and then dilating and eroding the image. This noise removal process accounted for frames distorted by passage of whiskers in front of the eye, and slight differences in face illumination between mice. For each session, appropriateness of fit was verified by overlaying the estimated pupil on the actual image for ∼20-50 randomly drawn frames. A single variable, the dark/light pixel thresholding value, could be changed to ensure optimal fitting for each session. Nose motion was extracted as the mean of pixel displacement in the ROI Y-axis estimated using an image registration algorithm (MATLAB ‘imregdemons’).

Whisker pad motion was estimated as the absolute difference in the whisker pad ROI between frames (MATLAB ‘imabsdiff’; this was sufficiently accurate to define whisking periods, and required much less computing time than ‘imregdemons’). Whisking was determined as the crossing of pad motion above a threshold, and whisking bouts were made continuous by convolving pad motion with a smoothing kernel. Licks were timestamped as the moment pixel intensity in the ROI in between the face and the lick port crossed a threshold.

Body movement was summarized as basket movements recorded by a triple-axis accelerometer (Adafruit, ADXL335) attached to the underside of a custom-designed 3D-printed basket suspended from springs (Century Spring Corp, ZZ3-36). Relative basket position was tracked by low-pass filtering accelerometer data at 2.5 Hz. Stimulations and cue deliveries were coordinated with custom-written software using Arduino Mega hardware (www.arduino.cc). All measurement and control signals were synchronously recorded and digitized (at 1 kHz for behavioral data, 10 kHz for fiber photometry data) with a Cerebus Signal Processor (Blackrock Microsystems). Data was analyzed using Matlab software (Mathworks).

### Preparatory and reactive measures and abstract learning trajectories

To describe the relationship between behavioral adaptations and reward collection performance, for each mouse in the control group a generalized linear model (GLM) was created to predict reward collection latency from preparatory and reactive predictor variables on each trial. Preparatory changes in licking, whisking, body movement, and pupil diameter were quantified by measuring the average of each of those signals during the 1 s delay period preceding cued rewards. The nose motion signal was not included as it did not display consistent preparatory changes.

Reactive responses in the whisking, nose motion, and body movement were measured as the latency to the first response following reward delivery. For whisking, this was simply the first moment of whisking following reward delivery. For nose motion, the raw signal was convolved with a smoothing kernel and then the first response was detected as a threshold crossing of the cumulative sum of the signal. For body movement, the response was detected as the first peak in the data following reward delivery. On occasional trials no event was detected within the analysis window. Additionally, discrete blocks of trials were lost due to data collection error for mouse 3-session 7, mouse4-session 5, and mouse9-session 4. In order to fit learning curves through these absent data points, missing trials were filled in using nearest neighbor interpolation.

Trial-by-trial reward collection latencies and predictor variables (preparatory licking, whisking, body movement, pupil diameter; and reactive nose motions, whisking, and body movement) were median filtered (MATLAB ‘medfilt1(signal,10)’) in order to minimize trial-to-trial variance in favor of variance due to learning across training. Collection latency was predicted from z-scored predictor variables using MATLAB ‘glmfit’ to fit β values for each predictor. The unique explained variance of each predictor was calculated as the difference in explained variance between the full model and a partial model in which β values were fit without using that predictor.

Preparatory and reactive predictor variables were used to define abstract learning trajectories which were plots of collection latency against the inferred reactive and preparatory variables for each of the first 800 cue-reward trials of training. Reactive and preparatory variables were calculated as the first principal component of the individual reactive and preparatory variables used in the GLM fits. For visualization we fit a parametric model to all 3 variables (single exponential for preparatory, double exponentials for reactive and latency using MATLAB ‘fit’ function). Quality of fits and choice of model were verified by visual inspection of all data for all mice. An individual mouse’s trajectory was then visualized by plotting downsampled versions of the fit functions for latency, reactive and preparatory. Arrowheads were placed at logarithmically spaced trials.

In order to quantify the total amount of preparatory behavior in each mouse at a given point in training (“final preparatory behavior”, Fig. 2e), each preparatory measure (pupil, licking, whisking, body movement) was z-scored and combined across mice into a single data matrix. The first principal component of this matrix was calculated and loading onto PC1 was defined as a measure of an inferred underlying ‘preparatory’ component of the behavioral policy. This created an equally weighted, variance-normalized combination of all preparatory measures to allow comparisons between individual mice. An analogous method was used to reduce the dimensionality of reactive variables down to a single ‘reactive’ dimension that captures the majority of variance in reactive behavioral variables across animals (Ext. Data Fig 2g). Initial NAc-DA signals were predicted from trained behavior at trials 700-800 by multiple regression (specifically, pseudoinverse of the data matrix of reactive and preparatory variables at the end of training multiplied by data matrix of physiological signals for all animals)(Ext. Data Fig. 2f).

### Combined fiber photometry and optogenetic stimulation

In the course of a single surgery session, DAT-Cre::ai32 mice received:

(1) Bilateral injections of AAV2/1-CAG-FLEX-jRCaMP1b in the VTA (150 nL at the coordinates -3.1 mm A/P, 1.3 mm M/L from bregma, at depths of 4.6 and 4.3 mm) or in the SNc (100 nL at the coordinates -3.2 mm A/P, 0.5 mm M/L, depth of 4.1, mm).
(2) Custom 0.39 NA, 200 μm fiber cannulas implanted bilaterally above the VTA (-3.2 mm A/P, 0.5 mm M/L, depth of -4.1 mm).
(3) Fiber cannula implanted unilaterally in the dorsomedial striatum (DS; 0.9 mm A/P, 1.5 mm M/L, depth of 2.5 mm) and nucleus accumbens core (NAc; 1.2 mm A/P, 0.85 mm M/L, depth of 4.3 mm). Hemisphere choice was counterbalanced across individuals. A detailed description of the methods has been published^53^.

Imaging began >20 days post-injections using custom-built fiber photometry systems (Fig. 2a)^53^. Two parallel excitation-emission channels through a 5-port filter cube (FMC5, Doric Lenses) allowed for simultaneous measurement of RCaMP1b and eYFP fluorescence, the latter channel having the purpose of controlling for the presence of movement artifacts. 470 nm and 565 nm fiber-coupled LEDs (M470F3, M565F3, Thorlabs) were connected to excitation ports with acceptance bandwidths of 465-490 nm and 555-570 nm respectively with 200 μm, 0.22 NA fibers (Doric Lenses). Light was conveyed between the sample port of the cube and the animal by a 200 μm core, 0.39 NA fiber (Doric Lenses) terminating in a ceramic ferrule that was connected to the implanted fiber cannula by a ceramic mating sleeve (ADAL1, Thorlabs) using index matching gel to improve coupling efficiency (G608N3, Thorlabs). Light collected from the sample fiber was measured at separate output ports (emission bandwidths 500-540 nm and 600-680 nm) by 600 μm core, 0.48 NA fibers (Doric Lenses) connected to silicon photoreceivers (2151, Newport).

A time-division multiplexing strategy was used in which LEDs were controlled at a frequency of 100 Hz (1 ms on, 10 ms off), offset from each other to avoid crosstalk between channels. A Y-cable split each LED output between the filter cube and a photodetector to measure output power. LED output power was 50-80 μW. This low power combined with the 10% duty cycle used for multiplexing, prevented local ChR2 excitation^53^ by 473 nm eYFP excitation. Excitation-specific signals were recovered in post-processing by only keeping data from each channel when its LED output power was high. Data was downsampled to 100 Hz, then band-pass filtered between 0.01 and 40 Hz with a 2nd-order Butterworth filter. Though movement artifacts were negligible when mice were head-fixed in the rig (the moveable basket was designed to minimize brain movement with respect to the skull^8^), according to standard procedure the least squares fit of the eYFP movement artifact signal was subtracted from the jRCaMP1b signal. dF/F was calculated by dividing the raw signal by a baseline defined as the polynomial trend (MATLAB ‘detrend’) across the entire session. This preserved local slow signal changes while correcting for photobleaching. Comparisons between mice were done using the z-scored dF/F.

Analysis windows were chosen to capture the extent of mean phasic activations following each kind of stimulus. For NAc-DA and VTA-DA, reward responses were quantified from 0-2 s after reward delivery and cue responses from 0-1 s after cue delivery. DS-DA exhibited significantly faster kinetics, and thus reward and cue responses were quantified from 0 to 0.75 s after delivery.

Somatic Chr2 excitation was performed with a 473 nm laser (50mW, OEM Laser Systems) coupled by a branching fiber patch cord (200 μm, Doric Lenses) to the VTA-implanted fibers using ceramic mating sleeves. 30 Hz burst activations (10 ms on, 23 ms off) were delivered with durations of either 150 ms for calibrated stimulation or 500 ms for large stimulations. For calibrated stimulation, laser power was set between 1-3 mW (steady state output) in order to produce a NAc-DA reactive of similar amplitude to the largest transients observed during the first several trials of the session. This was confirmed during analysis to have roughly doubled the size of reward-related NAc-DA transients (Fig. 5g). For large stimulations, steady state laser output was set to 10 mW.

### Computational learning model: ACTR

#### Behavioral plant

An important aspect of this modeling work was to create a generative agent model that would produce core aspects of reward-seeking behavior in mice. To this end we focused on licking, which in the context of this task is the unique aspect of behavior critical for reward collection. A reader may look at the function *dlRNN_Pcheck_transfer.m* within the software repository to appreciate the structure of the plant model. We describe the function of the plant briefly here. It is well known that during consumptive, repetitive licking mice exhibit preparatory periods of ∼7Hz licking. We modeled a simple fixed rate plant with an active, ‘lick’ state that emitted observed licks at a fixed time interval of 150 ms. The onset of this lick pattern relative to entry into the lick state was started at a variable phase of the interlock interval (average latency to lick init from transition into lick state ∼100ms). Stochastic transitions between ‘rest’ and ‘lick’ states were governed by forward and backward transition rates. The reverse transition rate was a constant that depended upon the presence of reward {5e-3 ms without reward, 5e-1 ms with reward}. This change in the backwards rate captured the average duration of consumptive licking bouts. The forward rate was governed by the scaled policy network output and a background tendency to transition to licking as a function of trial time (analogous to an exponential rising hazard function; *τ*=100ms). The output unit of the policy network was the sum of the RNN output unit (constrained {-1,1} by the tanh activation function) and a large reactive transient proportional to the sensory weight ({0,max_scale}) where max_scale was a free parameter generally bounded from 5 to 10 during initialization. This net output was scaled by *S*=0.02/ms to convert to a scaled transition rate in the policy output.

Behavior of the plant for a range of policies is illustrated in Sup. Fig XX. A large range of parameterizations were explored with qualitatively similar results. Chosen parameters were arrived at by scanning many different simulations and matching average initial and final latencies for cue-reward pairings across the population of animals. More complicated versions (high-pass filtered, non-linear scaling) of the transition from RNN output to transition rate can be explored in the provided function. However, all transformations were found to produce qualitatively similar results and thus the simplest (scalar) transformation was chosen for reported simulations for clarity of presentation.

#### Recurrent neural network (RNN)

As noted in the main text the RNN component of the model and the learning rules used for training drew upon inspiration from^49^ that itself drew upon inspiration variants of node perturbation methods^50^ and the classic policy optimization methods known as ‘REINFORCE’ rules^1,25^. Briefly,^49^ demonstrated that a relatively simple learning rule that computed a nonlinear function of the correlation between a change in input and change in output multiplied by the change in performance on the objective was sufficiently correlated with the analytic gradient to allow efficient training of the RNN. We implemented a few changes relative to this prior work. Below we delve into the learning rule as implemented here or a reader may examine the commented open source code to get further clarification as well. First, we describe the structure of the RNN and some core aspects of its function in the context of the model. The RNN was constructed largely as described in^49^ and was very comparable to the structure of a re-implementation of that model in^82^.

Although we explored a range of parameters governing RNN construction, many examples of which are shown in Sup. Fig. 2, the simulations shown in the main results come from a network with 50 units (N_u_=50; chosen for simulation efficiency, larger networks were explored extensively as well), densely connected (P_c_=0.9), spectral scaling to produce preparatory dynamics (***g***=1.3), a characteristic time constant (***τ***=25ms), and a standard tanh activation function for individual units. Initial internal weights of the network (W_ij_) were assigned according to the equation:

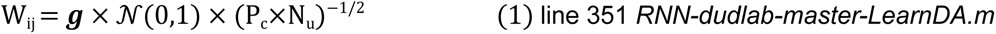

The RNN had a single primary output unit with activity that constituted the continuous time policy (*i.e. π(t)*) input to the behavior plant (see above), and a ‘feedback’ unit that did not project back into the network as would be standard, but rather was used to produce adaptive changes in the learning rate (described in more detail in “Learning rules” section below).

#### Objective function

Evaluation of model performance was calculated according to an objective function that defines the cost as the performance cost (2, “cost_P_”) and an optional network stability cost (3, “cost_N_”).

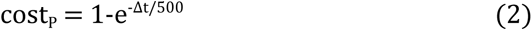

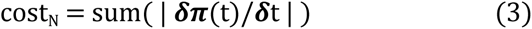

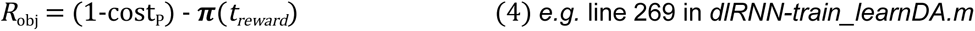

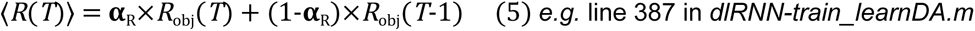

Where *T* is the trial index. In all presented simulations, 𝓌_N_=0.25. A filtered average cost, **R**, was computed as before^49^ with *α*_R_=0.75 and used in the update equation for changing network weights via the learning rule described below. For all constants a range of values were tried with qualitatively similar results. The performance objective was defined by cost_P_ where Δt is the latency to collected reward after it is available. The network stability cost (cost_N_) penalizes high-frequency oscillatory dynamics that can emerge in some (but not all) simulations. Such oscillations are inconsistent with observed dynamics of neural activity to date.

#### Identifying properties of RNN required for optimal performance

In order to examine what properties of the RNN were required for optimal performance, we scanned through thousands of simulated network configurations (random initializations of W_ij_) and ranked those networks according to their mean cost (R_obj_) when run through the behavior plant for 50 trials (an illustrative group of such simulations is shown in Sup. Fig. 2). This analysis revealed a few key aspects of the RNN required for optimality. First, a preparatory policy that spans time from the detection of the cue through the delivery of water reward minimizes latency cost. Second, while optimal RNNs are relatively indifferent to some parameters (*e.g.* P_c_) they tend to require a coupling coefficient (g) ≧1.2. This range of values for the coupling coefficient is known to determine the capacity of a RNN to develop preparatory dynamics^83^. Consistent with this interpretation we found that optimal policies were observed uniquely in RNNs with large leading eigenvalues (Sup. Fig. 2; *i.e.* long time constant dynamics^84^). These analyses define the optimal policy as one that requires preparatory dynamics of output unit activity that span the interval between the cue offset and reward delivery and further reveal that an RNN with long timescale dynamics is required to realize such a policy. Intuitively: preparatory anticipatory behavior, or “conditioned responding”, optimizes reward collection latency. If an agent is already licking when reward is delivered the latency to collect that reward is minimized.

#### RNN Initialization for simulations

All mice tested in our experiments began training with no preparatory licking to cues and a long latency (∼1 second or more) to collect water rewards. This indicates that animal behavior is consistent with an RNN initialization that has a policy *π*(t)∼0 for the entire trial. As noted above there are many random initializations of the RNN that can produce clear preparatory behavior and even optimal performance. Thus, we performed large searches of RNN initializations (random matrices W_ij_) and used only those that had ∼0 average activity in the output unit. We used a variety of different initializations across the simulations reported in Fig. 4 and indeed there can be substantial differences in the observed rate of convergence depending upon initial conditions (as there are across mice as well). For simulations of individual differences in Fig. 4h-j 6 distinct network initializations were chosen (as described above) and paired comparisons were made for the control initialization and an initialization in which the weights of the inputs from the reward to the internal RNN units were tripled.

#### Learning rules

Below we articulate how each aspect of the model acronym, ACTR (<u>A</u>daptive rate <u>C</u>ost of performance to <u>R</u>EINFORCE), is reflected in the learning rule that governs updates to the RNN. The connections between the variant of node perturbation used here and REINFORCE^25^ has been discussed in detail previously^49^. There are two key classes of weight changes governed by distinct learning rules within the ACTR model. First, we will discuss the learning that governs changes in the ‘internal’ weights of the RNN (W_ij_). The idea of the rule is to use perturbations (1-10Hz rate of perturbations in each unit; simulations reported used 3Hz) to drive fluctuations in activity and corresponding changes in the output unit that could improve or degrade performance. To solve the temporal credit assignment problem we used eligibility traces similar to those described previously^49^. One difference here was that the eligibility trace decayed exponentially with a time constant of 500 ms and it was unclear whether decay was a feature of prior work. The eligibility trace (ℯ) for a given connection *i,j* could be changed at any time point by computing a nonlinear function (𝒮) of the product of the derivative in the input from the *i*th unit (𝓍_i_) and the output rate of the *j*th unit (𝓇_j_) in the RNN according to the equation:

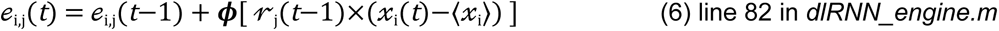

As noted by Miconi, the function 𝒮 need only be a signed, nonlinear function. Similarly, in our simulations we also found that a range of functions could all be used. Typically, we either used *ϕ(y)=y^3^* or *ϕ(y)=|y|*y* and simulations presented were generally the latter which runs more rapidly.

The change in a connection weight (W_ij_) in the RNN in the original formulation^49^ is then computed as the product of the eligibility trace and the change in performance error scaled by a learning rate parameter. Our implementation kept this core aspect of the computation, but several critical updates were made and will be described. First, since the eligibility trace is believed to be ‘read out’ into a plastic change in the synapse by a phasic burst of dopamine firing^79^. Thus, we chose to evaluate the eligibility at the time of the computed burst of DA firing estimated from the activity of the parallel feedback unit (see below for further details). Again, models that do not use this can also converge, but in general converge worse and less similarly to observed mice. The update equation is thus,

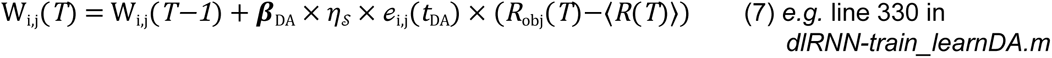

Where *η*_𝒮_is the baseline learning rate parameter and is generally used in the range {5e-4±1e-3} and *β*_DA_ is the ‘adaptive rate’ parameter that is a nonlinear function (sigmoid) of the sum of the derivative of the policy at the time of reward plus the magnitude of the reactive response component + T a tonic activity component (T=1 except in Fig. 1j where noted) and *ϕ* is a sigmoid function mapping inputs from {0,10} to {0,3} with parameters: **σ**=1.25, **μ**=7.

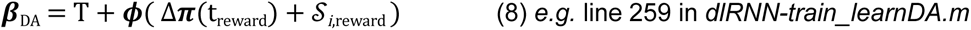

As noted in the description of the behavioral data described in Fig. 1 it is clear that animal behavior exhibits learning of both preparatory behavioral responses to the cue as well as reactive learning that reduces reaction times between sensory input (either cues or rewards) and motor outputs. This is particularly prominent in early training where a dramatic decrease in reward collection latency occurs even in the absence of particularly large changes in the preparatory component of behavior. We interpreted this reactive component as a ‘direct’ sensorimotor transformation consistent with the treatment of reaction times in the literature^85^ and thus reactive learning updates weights between sensory inputs and the output unit (one specific element of the RNN indexed as ‘o’ below). This reactive learning was also updated according to performance errors. In particular the difference between *R*_obj_(*T*) and the activity of the output unit at the time of reward delivery. For the cue updates were proportional to the difference between the derivative in the output unit activity at the cue and the performance error at the reward delivery. These rates were also scaled by the same *β*_DA_ adaptive learning rate parameter:

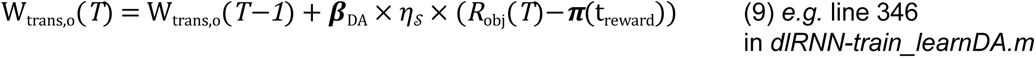

Where *η*_I_ is the baseline reactive learning rate and typical values were ∼0.02 in presented simulations (again a range of different initializations were tested).

We compared acquisition learning in the complete ACTR model to observed mouse behavior using a variety of approaches. For (1) we scanned ∼2 orders of magnitude for two critical parameters *η*_I_ and *η*_W_ (Supp Fig XX) and (2) aimed to sample the model across a range of initializations that approximately covered the range of learning curves exhibited by control mice. To scan this space we followed the following procedure. We initialized 500-1000 networks with random internal weights and initial sensory input weights (as described above). Since no mice that we observed initially exhibited sustained licking we selected 6 network initializations with preparatory policies approximately constant and 0. For these 6 net initializations we ran 24 simulations with 4 conditions for each init. Specifically, we simulated input vectors with initial weights 𝒮=[0.1, 0.125, 0.15, 0.175] and baseline learning rates *η*_I_ = [2, 2.25, 2.5, 2.75]*8e-3.

Representative curves of these simulations are shown in (Fig. 1j).

#### Visualizing the objective surface

In order to visualize the objective surface that governs learning we scanned a range of policies (combinations of reactive and preparatory components) passed through the behavior plant. The range of reactive components covered was [0:1.1] and preparatory was [-0.25:1]. This range corresponded to the space of all possible policy outputs realizable by the ACTR network. For each pair of values a policy was computed and passed through the behavior plant 50 times to get an estimate of the mean performance cost. These simulations were then fit using a third order 2d polynomials (analogous to the procedure used for experimental data) and visualized as a 3D surface.

In the case of experimental data the full distribution of individual trial data points across all mice (N=7200 observations) was used to fit a 3nd order, 2d polynomial (MATLAB; ‘fit’). Observed trajectories of preparatory vs reactive were superimposed on this surface by finding the nearest corresponding point on the fit 2d surface for the parametric preparatory and reactive trajectories. These data are presented in Fig. 1j.

#### Simulating closed-loop stimulation of mDA experiments

We sought to develop an experimental test of the model that was tractable (as opposed to inferring the unobserved policy for example). The experimenter in principle has access to real-time detection of licking during the cue-reward interval. In simulations this also can easily be observed by monitoring the output of the behavioral plant. Thus, in the model we kept track of individual trials and the number of licks produced in the cue-reward interval. For analysis experiments (Fig. 4h) we tracked these trials and separately calculated the predicted DA responses depending upon trial type classification. For simulations in Fig. 4h we ran simulations from the same initialization in 9 replicates (matched to the number of control mice) and error bars reflect the standard error.

To simulate calibrated stimulation of mDA neurons, we multiplied the adaptive rate parameter, *β*_DA_, by 2 based upon the same trial classification. Specifically, on trials with no licks detected (stim lick−) or on trials with at least one lick detected (stim lick+). For simulations reported in Fig. 4i we used 3 conditions: control, stim lick−, stim lick+. For each of these 3 conditions we ran 9 simulations (3 different initializations, 3 replicates) for 27 total learning simulations (800 trials). This choice was an attempt to estimate the expected experimental variance since trial classification scheme is an imperfect estimate of underlying policy.

#### Pseudocode summary of model

Here we provide a description of how the model functions in pseudocode to complement the graphical diagrams in the main figures and the discursive descriptions of individual elements that follow here in the Methods.

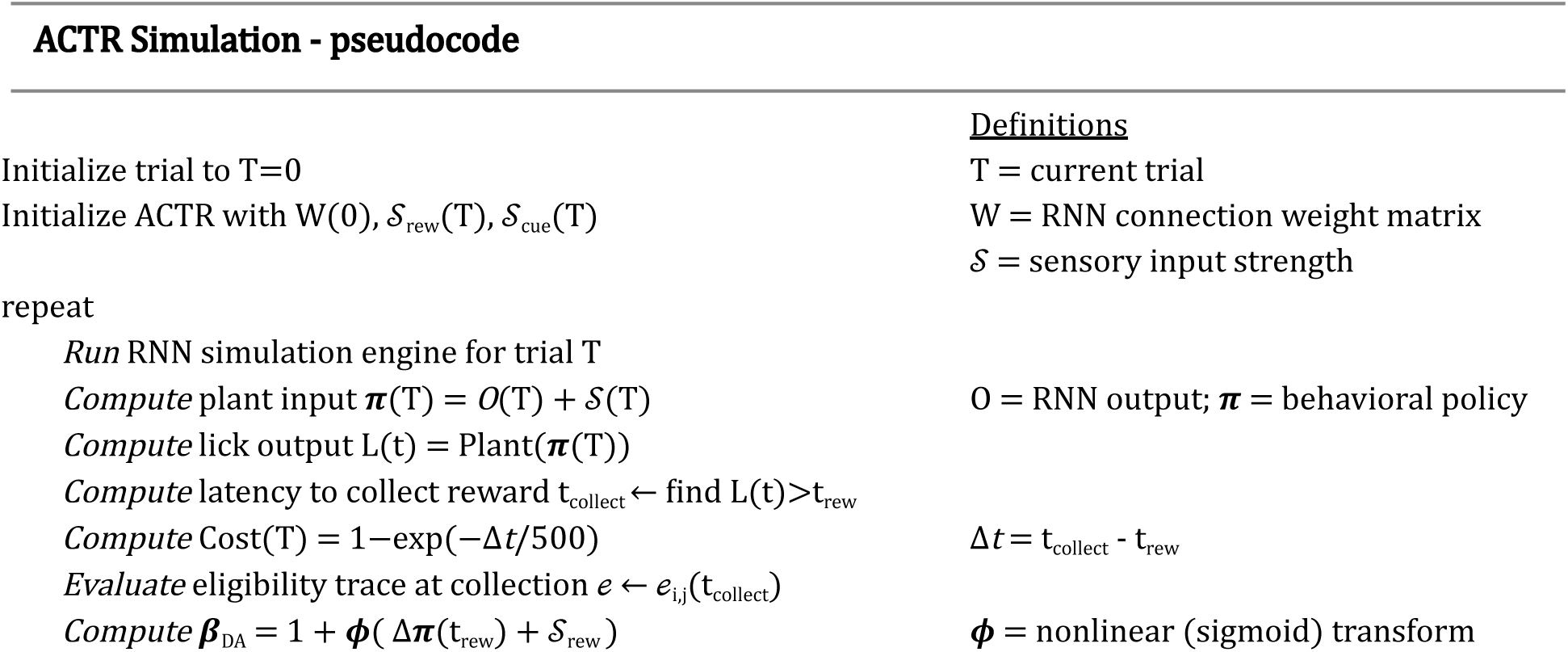
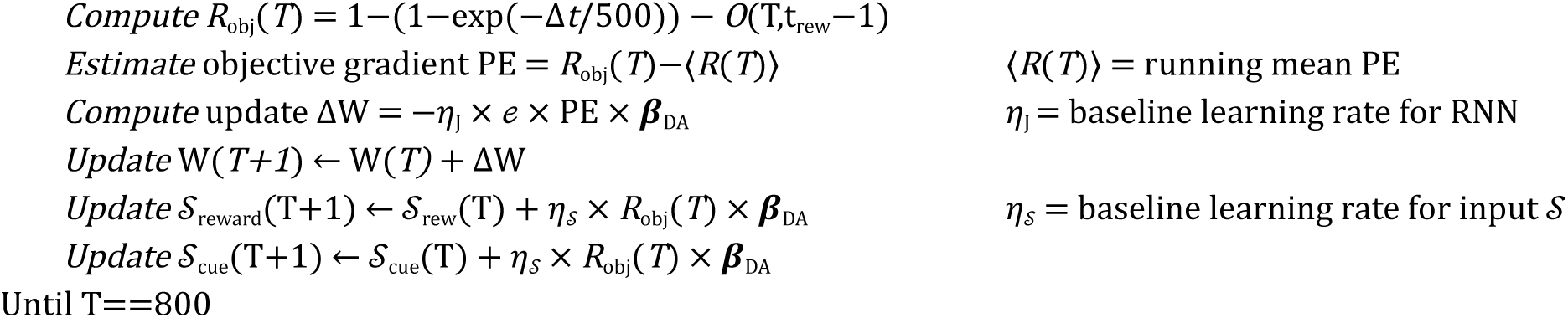

#### ACTR model variants

In Figure 1j we consider 3 model variants equivalent to DA signaling performance errors, DA depletion, and no phasic DA activity - all manipulations that have been published in the literature. To accomplish these simulations we (1) changed *β*_DA_ to equal PE; (2) changed *β*_DA_ offset to 0.1 from 1; (3) changed *β*_DA_ to equal 1 and removed the adaptive term.

In Figure 3 & 4 calibrated stimulation was modeled as setting *β*_DA_ to double the maximal possible magnitude of *β*_DA_ under normal learning. In Figure 3c-e, Figure 4h we modeled uncalibrated DA stimulation as setting PE=+1 in addition to the calibrated stimulation effect.

#### Temporal difference learning model

To model a standard temporal difference value learning model we reimplemented a previously published model that spanned a range of model parameterizations from^86^.

#### Code availability

All code relating to simulating the ACTR model and for a reader to explore both described parameterizations and explore a number of implemented, but unused in this manuscript, features can be found at https://github.com/dudmanj/RNN_learnDA. Specific line numbers are provided within the code for a subset of critical computations in the model.

#### Histology

Mice were killed by anesthetic overdose (isoflurane, >3%) and perfused with ice-cold phosphate-buffered saline (PBS), followed by paraformaldehyde (4% wt/vol in PBS). Brains were post-fixed for 2 h at 4° C and then rinsed in saline. Whole brains were then sectioned (100 μm thickness) using a vibrating microtome (VT-1200, Leica Microsystems). Fiber tip positions were estimated by referencing standard mouse brain coordinates^87^.

#### Statistical analysis

Two-sample, unpaired comparisons were made using Wilcoxon’s rank sum test (MATLAB ‘ranksum’); paired comparisons using Wilcoxon signed-rank test (MATLAB ‘signrank’). Multiple comparisons with repeated measures were made using Friedman’s test (MATLAB ‘friedman’). Comparisons between groups across training were made using 2-way ANOVA (MATLAB ‘anova2’). Correlations were quantified using Pearson’s correlation coefficient (MATLAB ‘corr’). Linear regression to estimate contribution of fiber position to variance in mDA reward signals was fit using MATLAB ‘fitlm’. Polynomial regression to fit objective surfaces were 3rd order and (MATLAB ‘fit’). Errors are reported as standard errors of the mean (s.e.m.). All sample sizes refer to the number of mice in the sample.

#### Data Availability

The data and custom code used to generate results supporting the findings of this study are within https://github.com/DudLab - including both modeling code (https://github.com/DudLab/RNN_learnDA) and analysis code (https://github.com/DudLab/TONIC).

**Extended Data Figure 1.**
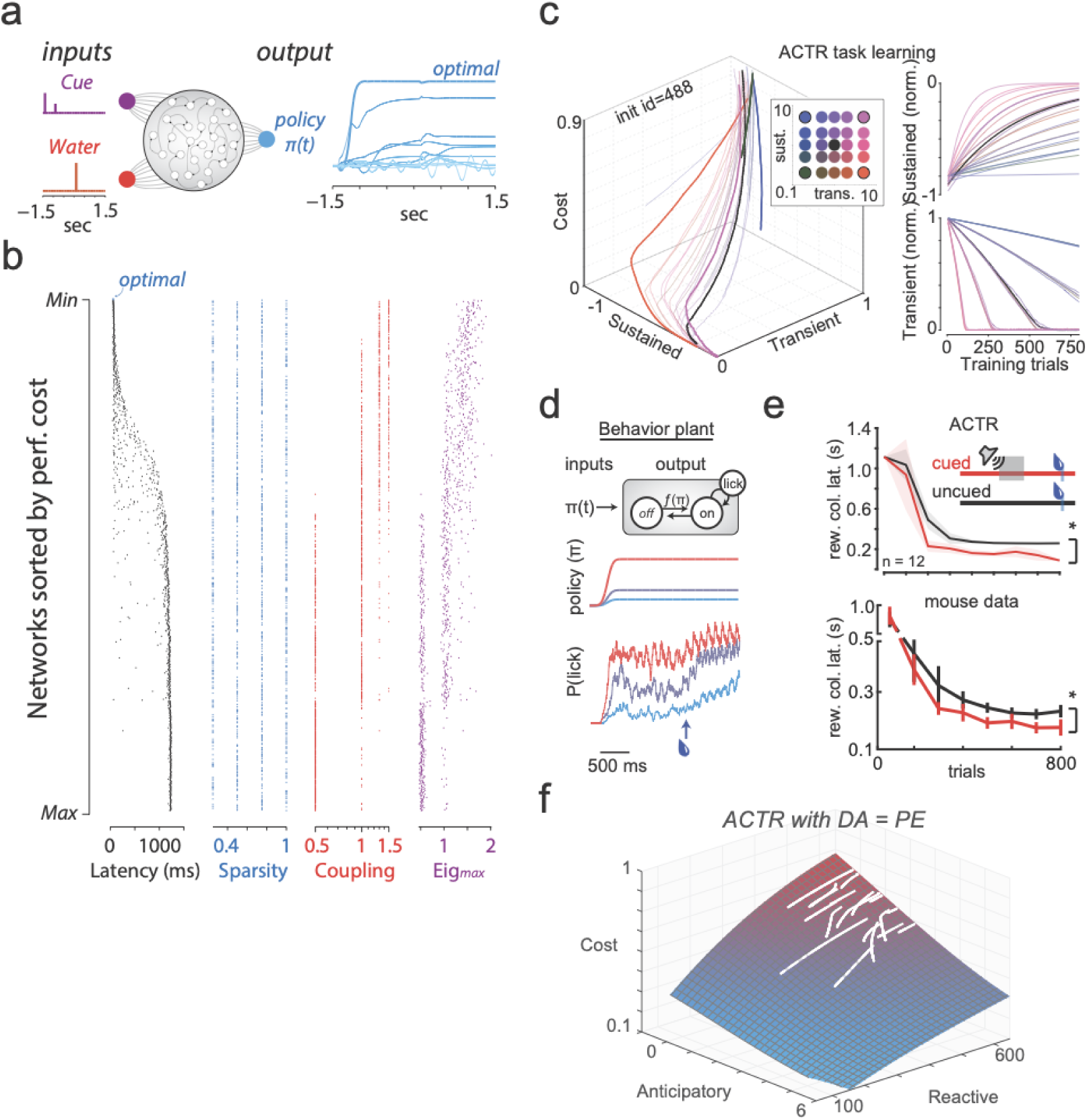
ACTR model details. **a)** top) Schematic of ACTR policy recurrent neural network (RNN) and licking output from example different network initialization (right) **b)** Thousands of randomized initial network configurations ranked according to their performance cost (Fig. 1h; cost is a combination of latency and a network variance cost, see methods for details). Displayed are the latency to collect reward (black), network sparsity (blue), coupling coefficient (red), leading eigenvalue (purple). This analysis reveals a few key aspects. First, a sustained policy that spans time from the detection of the cue through the delivery of water reward is necessary to minimize latency cost. Second, while optimal RNNs are relatively indifferent to some parameters (sparsity of connectivity) they tend to require a strong coupling coefficient which is known to determine the capacity of a RNN to develop sustained dynamics^83^, and thus optimal policies were observed uniquely in RNNs with large leading eigenvalues (i.e. long time constant dynamics^84^). These analyses indicate that there are realizable RNN configurations sufficient to produce an optimal policy, given an effective learning rule. **c)** Different ratios of sustained vs transient learning rates (inset color code) produced a range of trajectories similar to observed trajectories in individual mice (Fig. 1e). **d)** (top row) Licking behavior was modeled as a two state ({off,on}) plant that emitted 7 Hz lick bouts from the ‘on’ state. Forward transition rate (off→on) was determined by a policy *π(t)*. Reverse transition rate (on→off) was a constant modulated by the presence of water. Bottom three rows illustrate example licking behavior produced by the plant for three different constant policies (red, purple, blue) before and after water reward delivery (vertical black line) for 100 repetitions of each policy. **e)** Learning quantified by a decrease in reward collection latency over training. As training progressed, a predictive cue led to faster reward collection (red) as compared to uncued probe trials (black) in both the ACTR model (top, cued: 146 ± 21 ms, uncued: 205 ± 7 ms, p =0.01) and mouse data (bottom, cued: 176 ± 26 ms, uncued: 231 ± 23 ms, p =0.03) **f)** Cost surface (red = high, blue = low) overlaid with trajectories of individual initializations (white) as in Fig 1g, except having a constant learning rate and using the DA-like feedback unit to set the performance error (PE) instead of an adaptive learning rate (β), showing that the DA signal does not perform well as the PE in the ACTR learning rule

**Extended Data Figure 2.**
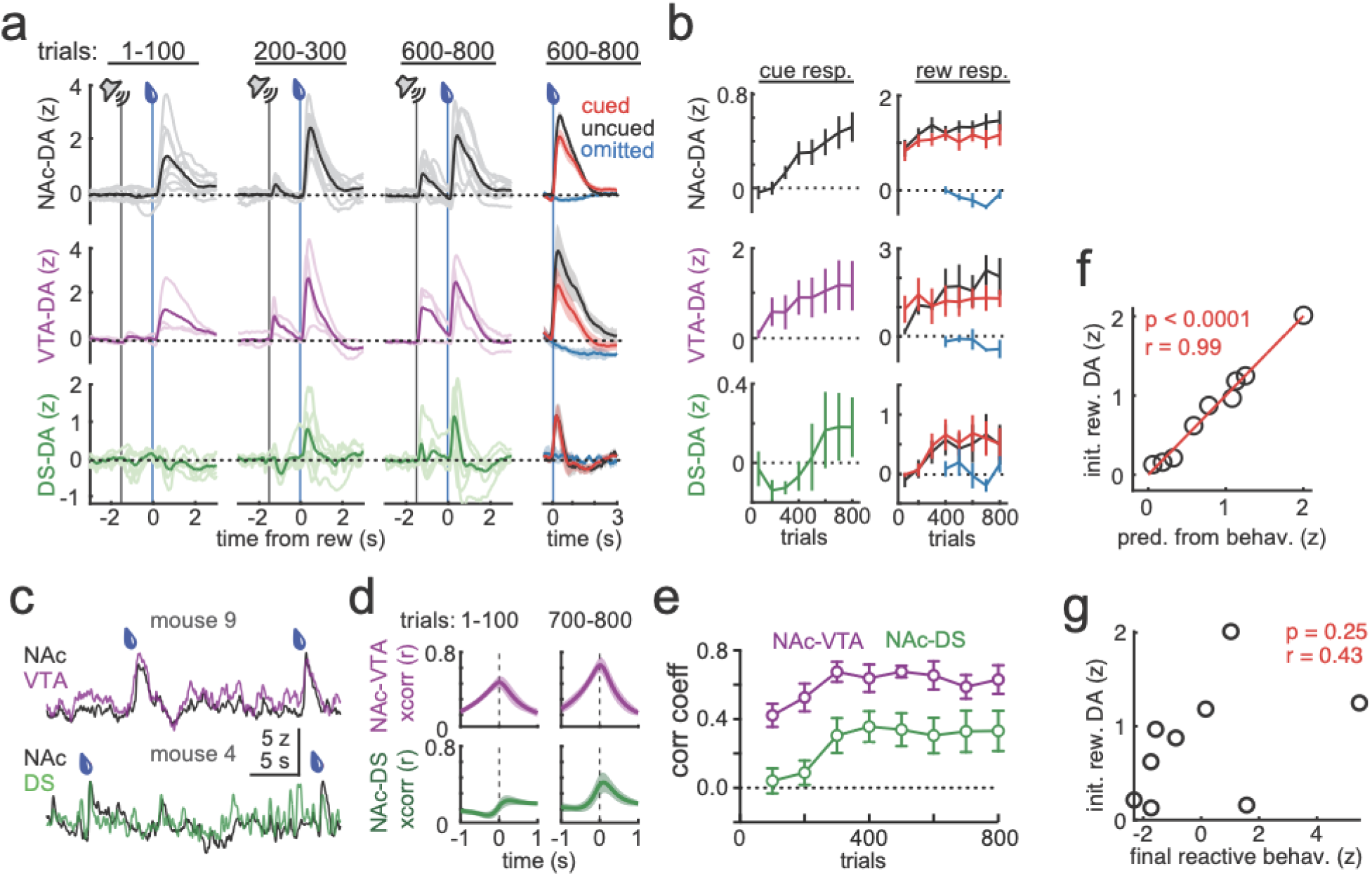
Dopamine signals across learning. **a)** (left 3 columns) jRCaMP1b DA signals in the nucleus accumbens core (NAc, black, n=9) and simultaneous recordings in the ventral tegmental area (VTA, purple, n = 3) and dorsal striatum (DS, green, n = 6), for cued reward trials in the trial bins indicated across training. (right) Reward or omission signals in NAc (top), VTA (middle), and DS (bottom) in trials 600-800, for cued (red), uncued (black), and cued but omitted (blue) trials. **b)** Mean signals during the 1 s following cue delivery (left) and 2 s following reward delivery (right) across training for each brain region from panel (C). **c)** Example simultaneous recordings from NAc-VTA (top) and NAc-DA (bottom). **d)** Mean cross correlations for simultaneously measured NAc-VTA signals (top row, n =3) and NAc-DS signals (bottom row, n=6) in trials 1-100 (left) and trials 700-800 (right) within trial periods (1 second before cue to 3 seconds after reward). **e)** Peak cross correlation coefficients between NAc-VTA and NAc-DS signal pairs across training, within trial periods. **f)** Correlation of initial NAc-DA reward signals predicted from behavior measures in trials 700-800 (see Methods) to observed initial NAc-DA reward signals. **g)** No correlation of initial NAc-DA reward signals with final combined reactive behaviors

**Extended Data Figure 3.**
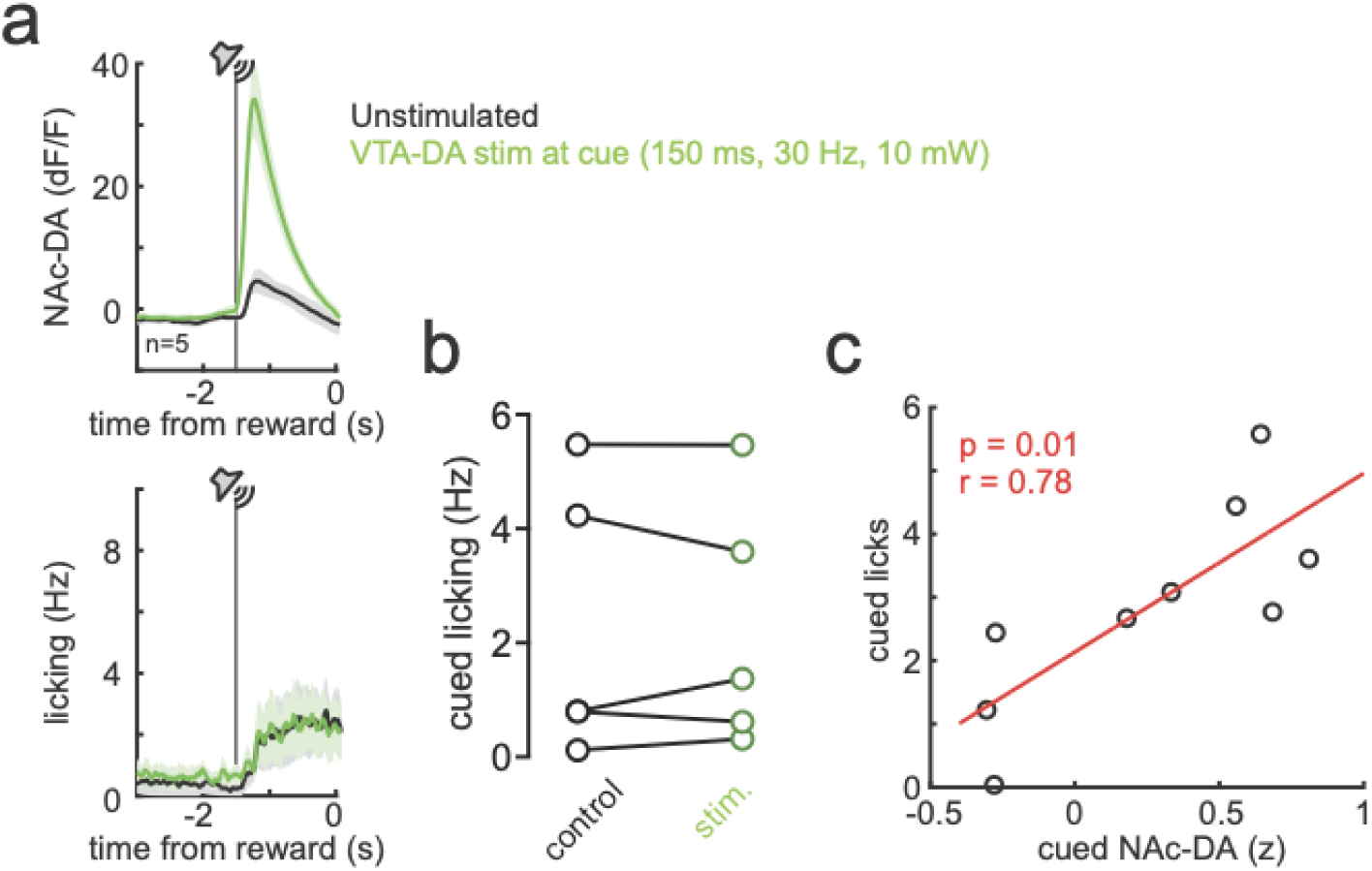
Augmenting mesolimbic cue signals does not affect cued behavior. **a)** To test for a causal connection between the size of mesolimbic DA cue responses and cued behavior, in a new session after regular training was complete, we delivered large, uncalibrated VTA-DA stimulation on a random subset of cued reward trials (light green). Shown are NAc-DA responses (top) and licking (bottom) for this session. **b)** Quantification of cued preparatory licking during the delay period for unstimulated (black) vs stimulated (green) trials. **c)** Cued licking was correlated with the size of NAc-DA cue responses across animals, even though manipulations did not support a causal relationship.

**Extended Data Figure 4.**
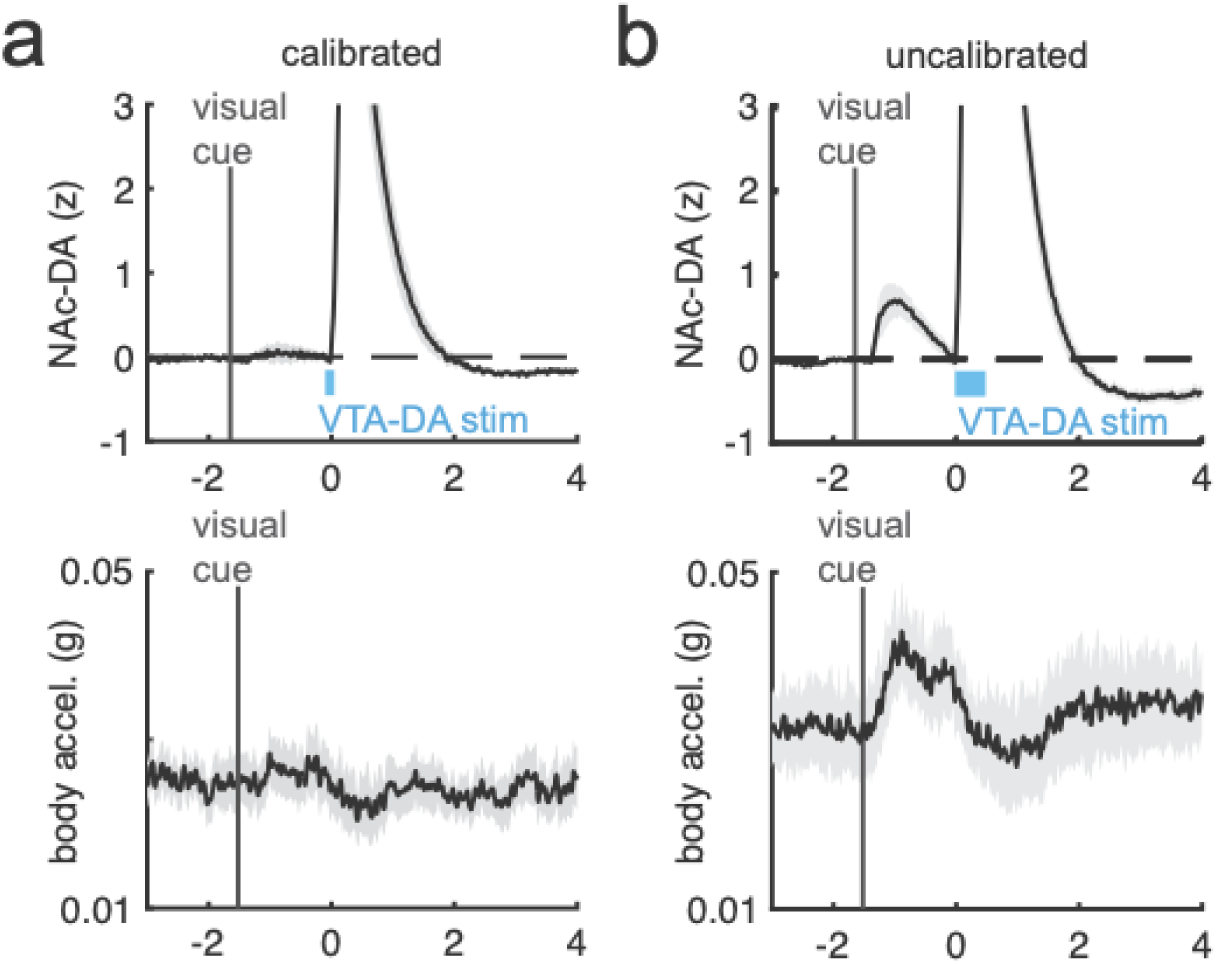
Movement correlates after learning induced by exogenous VTA-DA stimulation. **a)** NAc-DA responses (top) and body movement measured as acceleration of the basket holding the mice (bottom) at the end of training in the paradigm in Fig 3a-e in which a visual cue predicted VTA-DA stimulation calibrated to measured uncued reward responses **b)** Same as (a) except for larger, uncalibrated stimulation

**Extended Data Figure 5.**
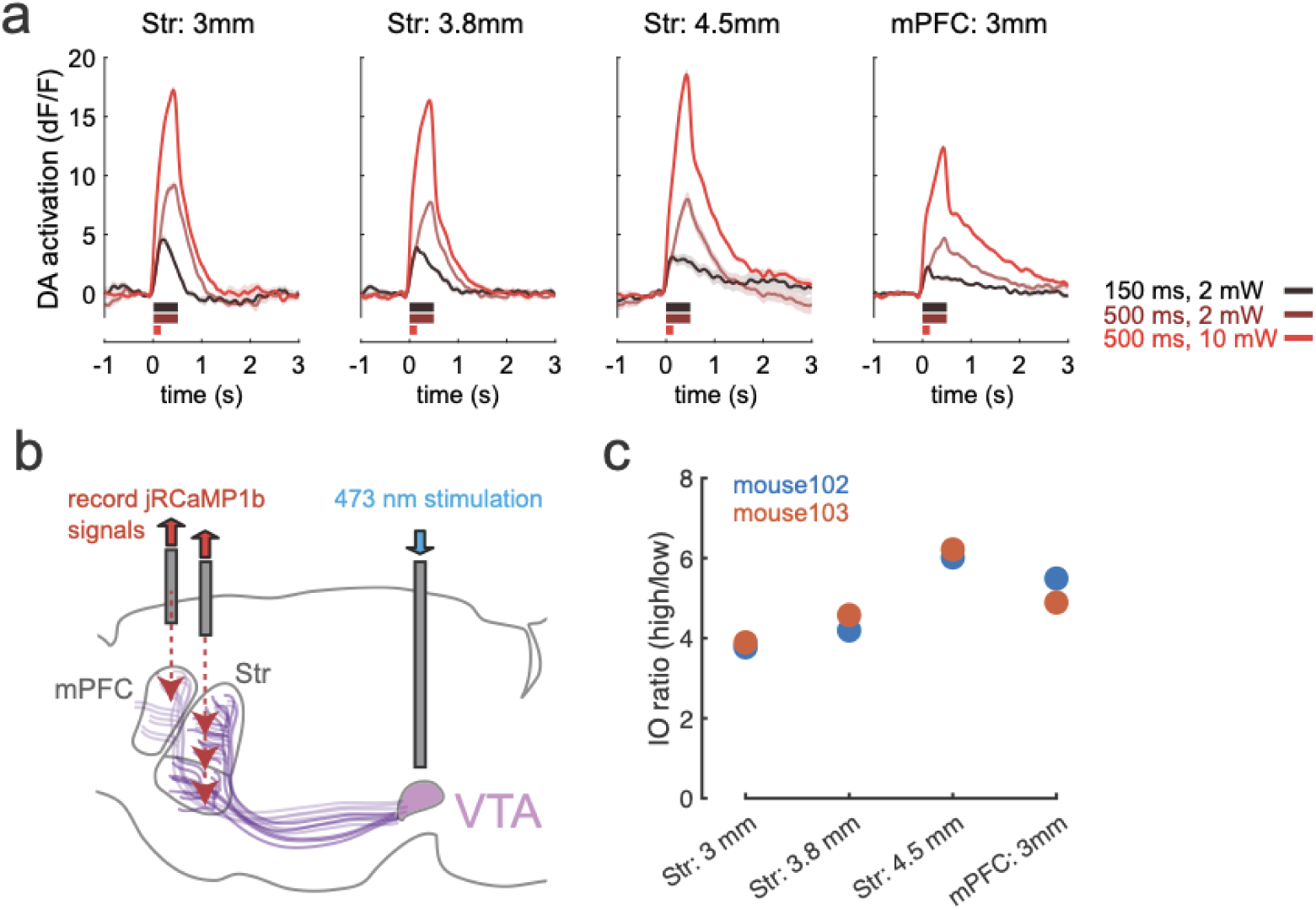
Exogenous stimulation produces similar input-output across sampled DA projection targets. **a)** DA responses measured through the same fiber for the low (black), medium (dark red), and high (bright red) stim parameters indicated at right, inserted either in the mPFC or at the indicated depth in the striatum **b)** Schematic of experiment **c)** Input-output ratio (response to high stim parameters divided by response to low stim parameters at each recording site in two separate mice (blue and orange)

**Extended Data Figure 6.**
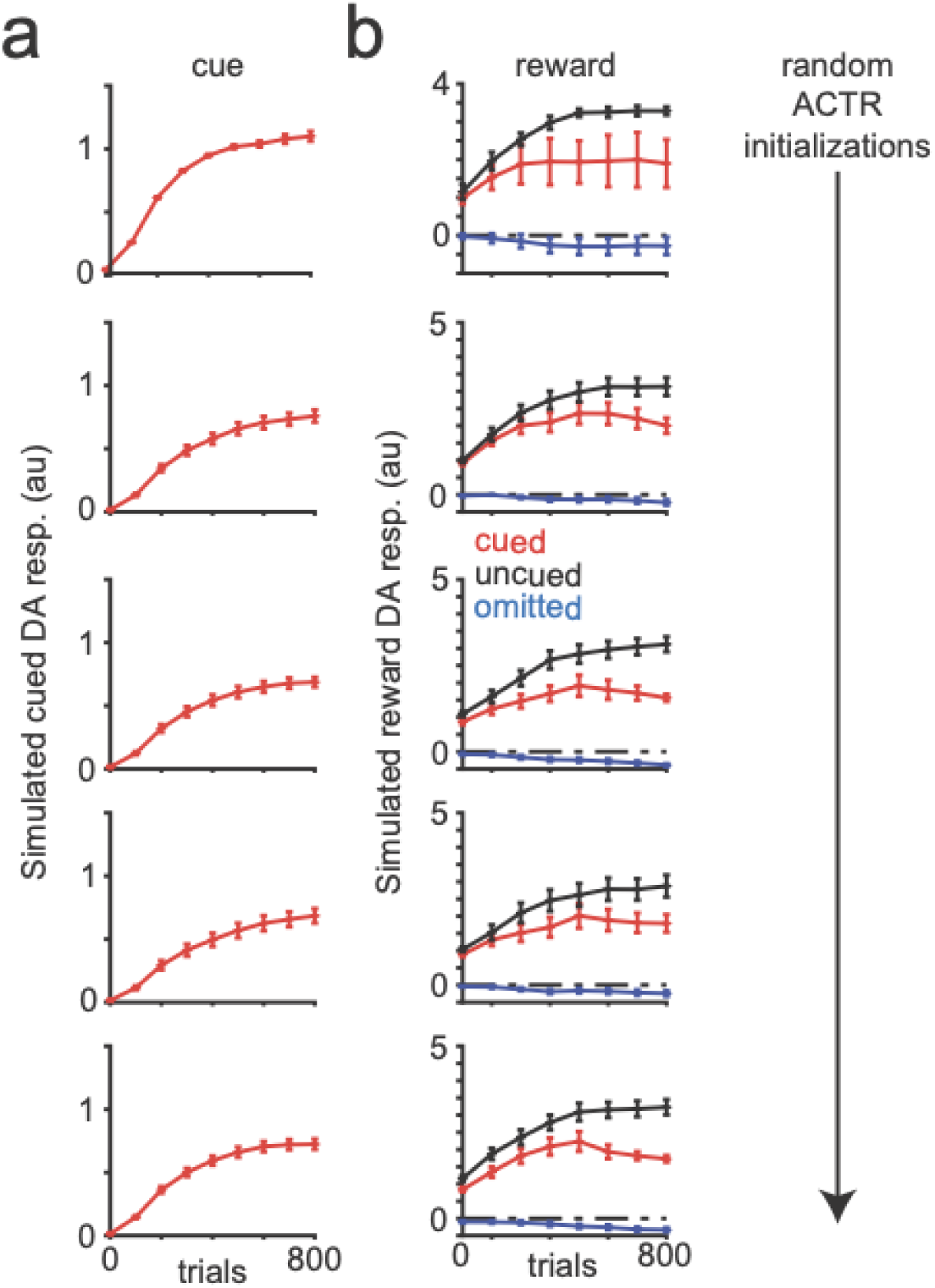
ACTR-modeled DA dynamics robust to initialization conditions. **a)** Predicted DA responses to cues for 5 random initializations of the ACTR model **b)** Predicted DA responses to rewards over the same initializations as (a), comparing cued (red), uncued (black) and omitted (blue) reward trials. RPE correlates emerged robustly across initializations

**Extended Data Figure 7.**
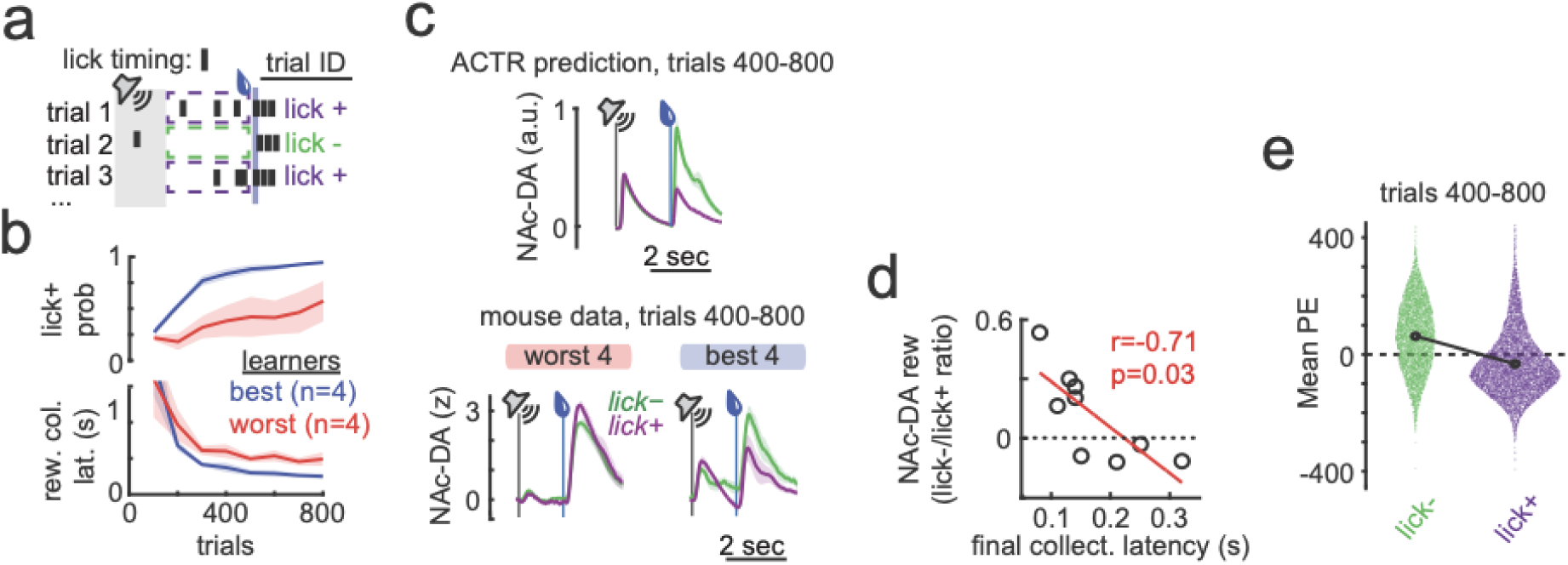
Relationship of NAc-DA signals to presence of preparatory licking. **a)** Trial types defined as “lick+” trials (purple) with at least one lick during the delay between cue and reward and “lick-” trials (green) with no delay period licking. **b)** Percent of total trials that are “lick+” (top) and reward collection latency (bottom), for best 4 (green) and worst 4 (purple) performing mice, as determined by their reward collection latency in trials 700-800. **c)** NAc-DA signals in the second half of training (trials 400-800) lick- (green) and lick+ (purple) trials, for ACTR simulations (top) and worst (left) and best (right) top 4 performing mice in terms of reward collection latency in trials 700-800. **d)** The ratio of NAc-DA reward signals on lick- vs lick+ trials was correlated with the final reward collection latency. **e)** The performance error (PE) can be computed for every trial. For trials 400-800 across 9 control mice, the distribution of PEs for lick+ (purple) and lick- (green) trials are shown as a swarmplot. The mean PE for each trial type is indicated by the connected black points; note the shift in mean from positive to negative.

**Extended Data Figure 8.**
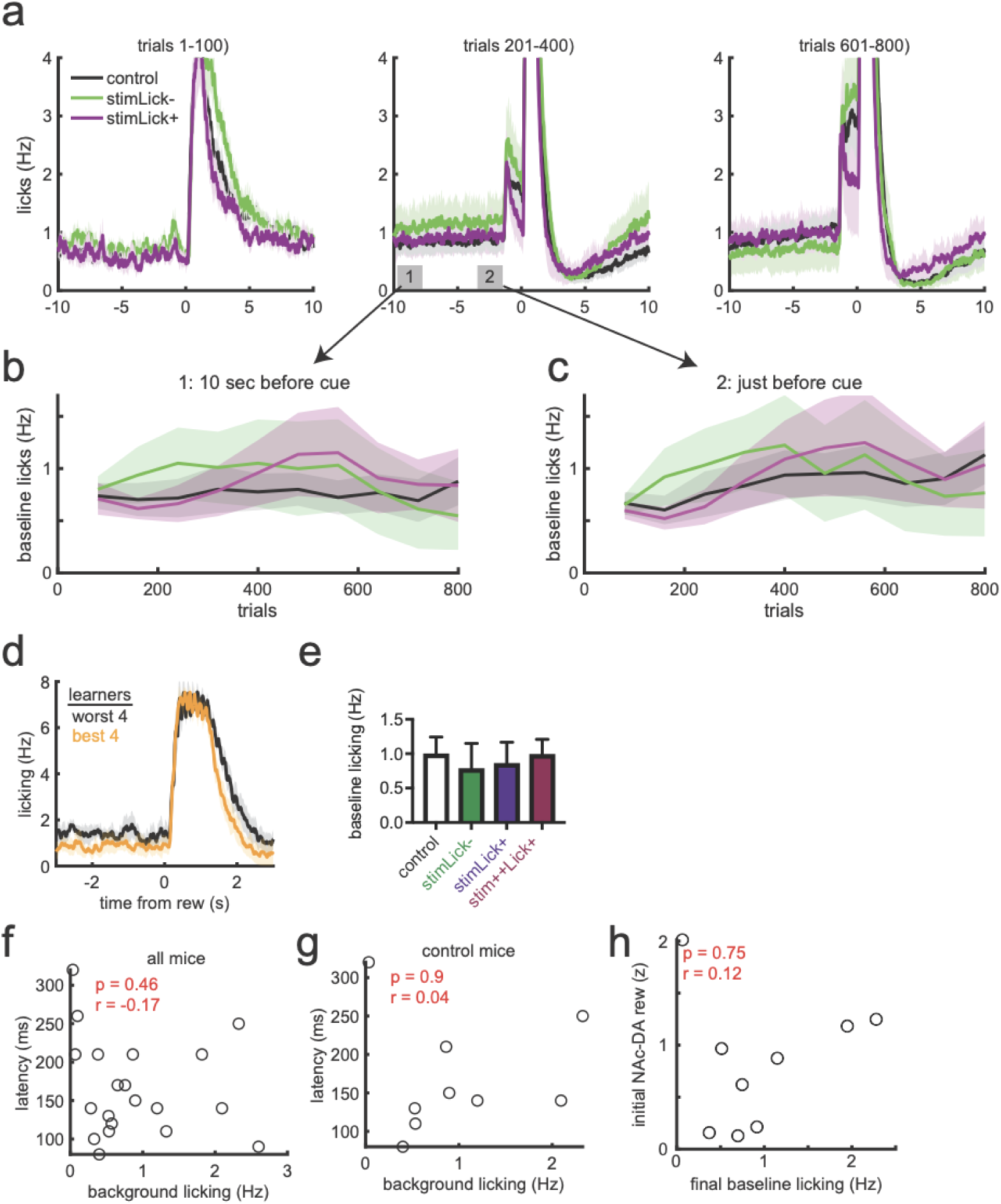
Baseline licking across training in all animals. **a)** Licking behavior showed in extended time before and after reward delivery to illustrate baseline intertrial licking behavior across the indicated training trials, for control (black), stimLick- (green), and stimLick+ (purple) mice **b)** Quantification of baseline licking at analysis epoch 1 (indicated in the middle trace of panel (a)) across training **c)** Quantification of baseline licking at analysis epoch 2 (indicated in the middle trace of panel (a)) across training **d)** Licking behavior over the 3 seconds preceding uncued trials at the end of training (trials 600-800) for the best 4 and worst 4 performing mice displayed an insignificant trend towards more baseline licking in bad learners **e)** Comparable baseline licking rate for all the experimental groups shown in Fig 1-4 **f)** No correlation between baseline licking and final latency to collect reward (a measure of learned performance) for all mice **g)** No correlation between baseline licking and final latency to collect reward (a measure of learned performance) for only control mice (Fig 1-2) **h)** No correlation between baseline licking and initial NAc-DA reward responses for control mice

**Extended Data Figure 9.**
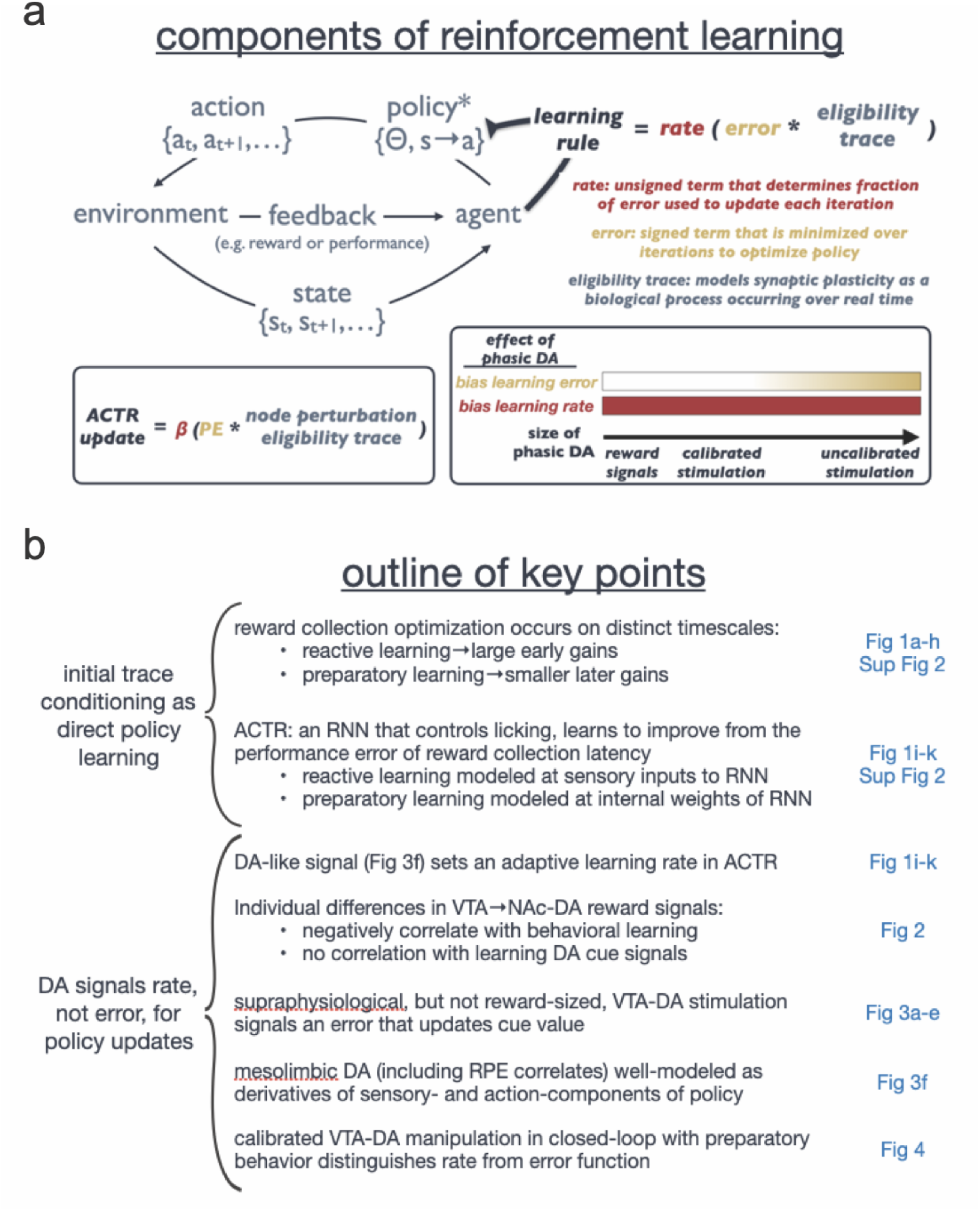
Logic Outline. **a)** In Reinforcement Learning, an agent learns iteratively from environmental feedback to improve a policy, which is a set of parameters (Θ) describing an action (a) that is performed given a state (s). In policy learning, the agent applies a learning rule **b)** Key points of the paper grouped by theme (left), with location in figures for primary supporting data (blue)

